# Ecological stability masks rapid, convergent evolution within the honey bee worker microbiome

**DOI:** 10.64898/2026.09.24.754198

**Authors:** Chris R. P. Robinson, Adam G. Dolezal, Irene L. G. Newton

## Abstract

While ecological dynamics within host-associated microbiomes are well-documented, how evolutionary processes unfold within them remains poorly understood. Using longitudinal metagenomics, we quantified ecological and evolutionary dynamics across over 1,000 bacterial genomes from honey bee worker microbiomes sampled across 12 colonies. We show that microbial community composition is strongly tied to colony identity and diverges continuously over time, rather than through discrete seasonal shifts. Despite this ecological turnover, most bacterial populations are stable, harboring extensive genetic variation shaped by strong purifying selection. Against this constrained backdrop, we identify thousands of rapid shifts in allele frequency that cannot be explained by strain replacement, population abundance, or genetic drift. Nonsynonymous variants exhibit localized signatures of selection at neighboring synonymous sites and independently recur within homologous proteins shared by phylogentically distant taxa found across independent colonies. Altogether, our findings demonstrate that ecological persistence can mask rapid evolutionary dynamics, revealing recurrent genetic signatures consistent with adaptive evolution across diverse bacteria.

## 2 Introduction

Host-associated microbiomes are complex ecosystems that harbor enormous potential for within-host bacterial evolution and adaptation. However, while ecological processes have been extensively studied across many model systems [1–16], we know far less about the evolutionary processes acting within populations of host-associated bacteria or among populations distributed across multiple hosts. Recent work has shown that microbial populations can evolve over ecologically relevant timescales [17–24], even in the absence of antibiotics. However, efforts to identify strong metagenomic signatures of adaptive evolution within host-associated microbiomes are complicated by these overlapping processes, such as changes in community composition or genetic drift.

The European honey bee (Apis mellifera) is an emerging model for studying population genetic processes within microbiomes due to the relative taxonomic simplicity, temporal stability, and metabolic specialization of its gut microbial community [25–29]. Worker-associated communities are dominated by a small number of core bacterial genera that remain comparatively stable within colonies, yet recent analyses have revealed substantial strain diversity within colonies and individual workers [25, 30–33]. Studies in this system have begun to resolve how priority effects [10, 34, 35], spatial partitioning [27, 36], interspecific interactions [26, 37], and seasonal variation structure these communities [38–40] over time. However, how these ecological processes structure genetic variation within bee-associated bacterial populations remains poorly understood.

Here, we use deeply sequenced longitudinal metagenomics from 12 honey bee colonies across three apiaries to resolve ecological and population-genetic dynamics over approximately one year. Community composition changed continuously and partly synchronously among colonies, while hundreds of bacterial populations persisted with stable abundance and extensive standing variation shaped predominantly by purifying selection. Against this constrained background, we identify thousands of rapid allele-frequency shifts, many occurring without major changes in population abundance or resident strain composition. We show that nonsynonymous shifts coincide with spatially and temporally localized change at neighboring synonymous sites and independently recur within homologous proteins across phylogenetically disparate taxa. Together, these signatures narrow thousands of rapid shifts to a conservative set of recurrent functional targets consistent with convergent adaptation in the honey bee gut microbiome

## 3 Results

### 3.1 Longitudinal metagenomics resolves diverse bacterial populations within the honey bee gut microbiome

To comprehensively characterize worker-associated microbial variation across space and time, we collected over 12,000 in-hive bees and larvae from 12 colonies across three apiaries and generated 298 metagenomic libraries (Figure 1A; see Methods). Colonies were sampled monthly for up to 12 months or until colony death (Figure 1B). Triplicate worker samples from each timepoint were co-assembled, generating 1,698 MAGs, of which 1,027 passed quality filters and dereplicated into 633 representative MAG clusters (rMAGs) at 95% ANI and 85% breadth. Dereplication patterns varied substantially among bacterial genera, reflecting broad differences in genome-level diversity within the worker microbiome (Extended Data 1, Supplemental Figures 1-2 for assembly and dereplication patterns (also see Supplemental Tables 1-2)).

**Figure 1.**
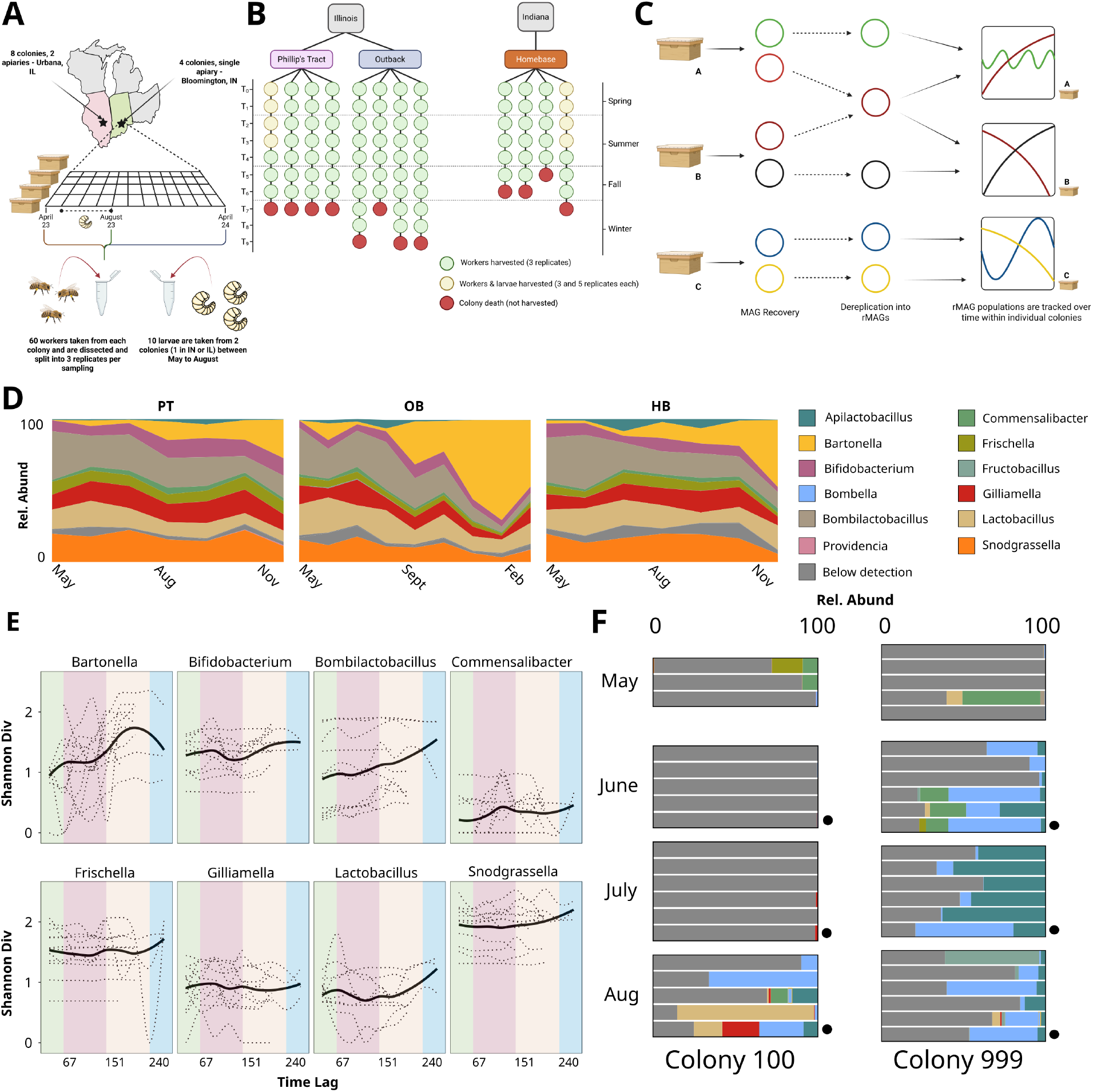
**A)** Experimental design and sampling strategy. Replicates refer to individual biological replicates taken at each colony and timepoint; individuals refer to individual colonies; populations refer to apiaries; metapopulations refer to groups of apiaries (same-state). **B)** Sampling timeline for each colony. Colony death references the month where colony populations had collapsed and therefore could not be sampled. Each timepoint refers to a separate month. **C)** Relative abundance (y-axis) throughout sampling (x-axis) within each of the three apiaries. Colors are representative of distinct genera. Coverage attributed to MAGs that failed detection thresholds are indicated in dark gray (Below detection). **D)** Shannon diversity (y-axis) for each genus associated with workers throughout the timecourse (x-axis). Time Lag references the number of days since the initial timepoint. Dotted lines represent individual colony trajectories while the solid black line represents an averaged trend. Colors within each square correspond to seasons (green: spring; red: summer; yellow: autumn; winter: fall.) **E)** Relative abundance for larval samples. Larvae are split into groups based on colony origin (x-axis) as well as timepoint (y-axis). Genera are demarcated by colors (using the same scheme as **A)**. Each individual bar is a single larvae, unless indicated by a black circle to the right of the bar. These samples consist of pools of 5 larvae.

We next investigated the abundance and distribution of recovered microbial populations across individual colonies using colony-specific rMAG databases for read recruitment (see Methods). We define population as any unique *rMAG x Colony* combination to account for rMAGs co-occurring across multiple individual colonies. Overall, populations were highly localized to a single colony, with only a single rMAG (*Commensalibacter* MAG 411) detected in all 12 colonies across all sampled timepoints. On average, each colony was stably associated with approximately 40 rMAGs (Supplemental Table 2 and Supplemental Figure 3), while genus-level relative abundance was comparatively similar across colonies and apiaries (Figure 1C for apiary abundance and Extended Data 2 and Supplemental Table 3). In contrast, larval samples exhibited weak microbial association. Most rMAGs failed our detection threshold, though recovered bacteria were most frequently associated with *Bombella* and *Apilactobacillus* (Figure 1D and Extended Data 2). Because of the paucity and inconsistency of bacterial populations associated with larvae (see Extended Data 3), we focused subsequent ecological and population genetic analyses on worker microbiomes. Within workers, Shannon diversity varied over time and among genera, generally increasing through Fall before declining in Winter (Figure 1E), although no individual population or genus showed consistent seasonal changes in abundance. In lieu of large changes of abundance of individual populations, we asked whether community dynamics were better explained by continuous compositional turnover within bee-associated microbial communities.

### 3.2 Measures of temporal community dissimilarity are similar among colonies and driven by few microbial populations

Having observed differences in the distribution of species diversity among genera and seasonal categories, we sought to better understand how patterns of species diversity were structured with respect to the hierarchical nature of our sampling and more nuanced measures of seasonal variation. Honey bee gut community composition was strongly influenced by colony identity (PERMANOVA: (*R*^2^ = 43.3%, *P <* 0.001). However, because reads from each colony were recruited against colony-specific rMAG databases, rMAG abundance was not treated as a common feature across colonies. Instead, corrected rMAG relative abundances were aggregated across genera for cross-colony analyses while biological replicate pools were retained as separate observations. After conditioning Bray-Curtis ordination on colony identity, residual community composition remained strongly organized across increasing time lag (Figure 2A, see Extended Data 4 for unconditioned NMDS). Consistent with this finding, we observed that Bray Curtis dissimilarity-lag relationships were significant and positive across colonies, with slopes *>* 0.3 in 10/12 colonies (Figure 2B).

**Figure 2.**
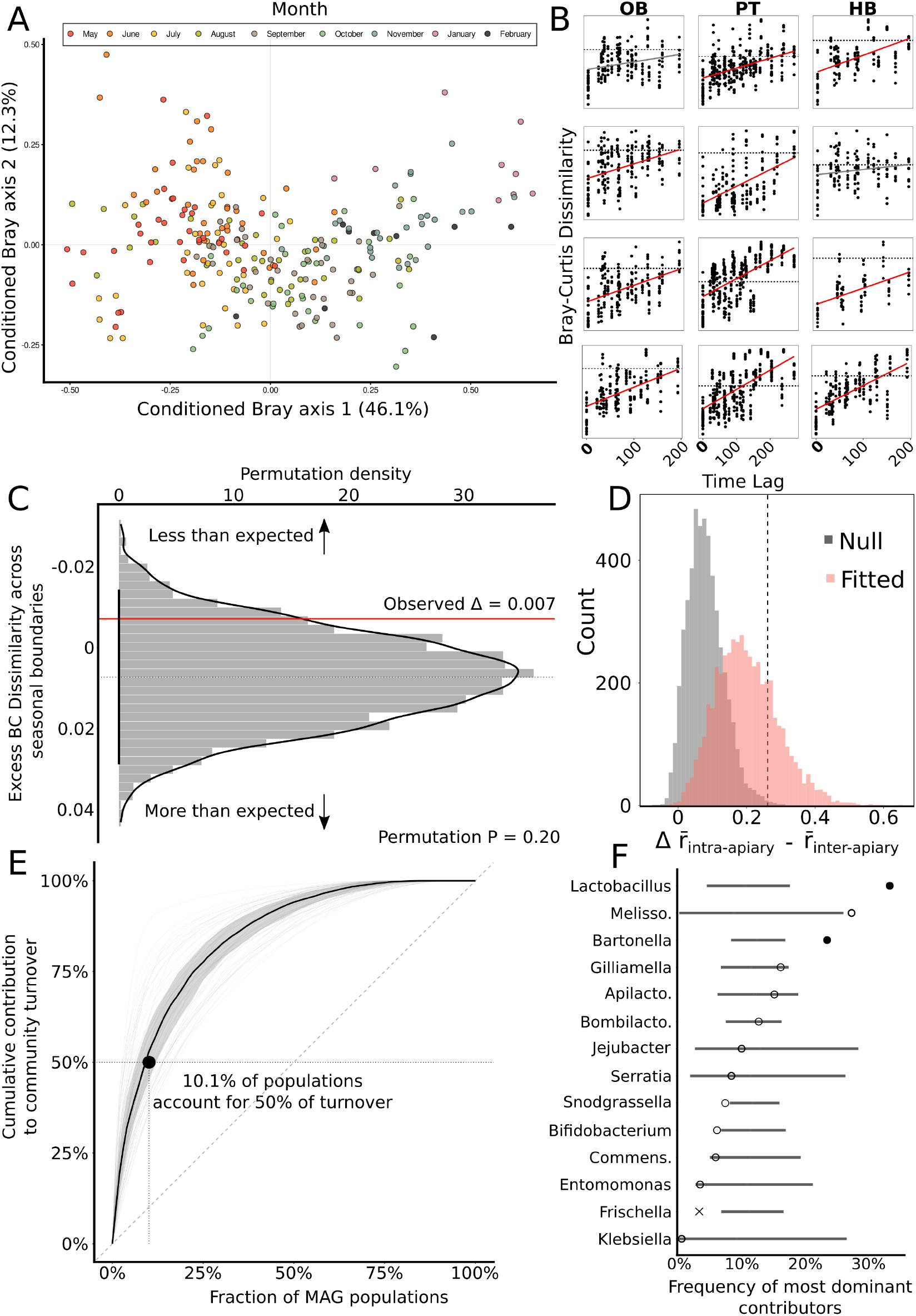
**A)** NMDS ordination of species abundances across metagenomes, colored by month. **B)** Bray-Curtis dissimilarity (y-axis) over time (x-axis) for individual colonies (squares), grouped by apiary (columns). Red lines indicate statistically significant increases in dissimilarity over time. **C)** Distribution of excess Bray-Curtis dissimilarity (y-axis) controlling for seasonal category and time lag. Red line denotes observed excess dissimilarity. **D)** Distributions of synchrony scores, calculated by subtracting inter-apiary colony scores from intra-apiary scores across bootstrap resamples. Black bar indicates the 95% bootstrap confidence interval. **E)** Lorenz curve depicting the fraction of MAG populations (x-axis) driving cumulative community turnover (y-axis). Permuted null trajectories are shown in gray, median contribution in black, and the black dot highlights the MAG fraction driving 50% of turnover. **F)** Contribution of dominant populations (x-axis) to community turnover across genera (y-axis). Filled circles indicate significant positive contributions, open circles denote non-significant associations, and ‘X’ marks genera contributing significantly less turnover than expected.

While increasing dissimilarity aligns with previously reported seasonal dynamics [38–40], Time Lag, Month, and Season capture the same temporal trajectory. We asked whether discrete seasonal categories contributed more dissimilarity than expected from time lag alone. Discrete seasonal categories contributed minimal excess dissimilarity beyond time lag alone Δ_*season*_ = 0.0071 bootstrap 95% CI, *−*0.034 to 0.01777) and did not exceed a within-colony circular-shift null (*P* = 0.196, Figure 2C). Thus, community turnover occurs primarily across continuous time rather than discrete seasonal transitions.

Although community composition diverged progressively through time, the magnitude of change varied among consecutive samplings. We therefore asked whether colonies tended to experience relatively large or small ecological shifts during the same sampling interval. We quantified synchrony between colony pairs as the Spearman correlation between their consecutive Bray-Curtis turnover trajectories (see Methods). Synchrony was stronger among colonies housed within the same apiary than among colonies from different apiaries (Figure 2D; 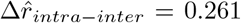, *P* = 0.003). This finding suggests that while individual colonies share a common temporal response trajectory with respect to compositional dissimilarity, deviations around this trajectory are more likely to be shared by colonies nearer to each other compared to more distant colonies.

Finally, we asked whether this turnover reflected homogenous change across the community or was concentrated among few taxa. Within each colony and interval, rMAG abundance change was expressed as its proportional contribution to total community turnover. A median of only 10.1% of rMAG populations accounted for 50% of total compositional change (Figure 2E), with populations belonging to *Lactobacillus* and *Bartonella* frequently observed as the strongest contributors (Figure 2F). These two genera also exhibited significantly high intra-genus Bray-curtis dissimilarity (see Extended Data 5). Together, our findings show that colony-specific honey bee worker gut microbiomes undergo continuous and partially synchronized ecological change through time, but that much of this turnover is driven by a comparatively small subset of bacterial populations.

### 3.3 Persistent microbial populations harbor extensive standing genetic variation

Having established strong patterns of population persistence and stability, we next explored patterns of subspecies diversity. Deep metagenomic sequencing allowed us to monitor SNVs (single nucleotide variants) within 497 populations that passed our prevalence and gene filters (see Methods). Consistent with previous work in the human gut microbiome [17, 19, 41–43], we found that standing genetic diversity varied tremendously among populations (Figure 3B and Supplemental Table 3). 61.7% were associated with at least 10^4^ SNVs within a given month while 38.1% associated with *<* 10^4^. Several populations maintained exceptionally high SNV burdens throughout sampling, whereas others remained persistently depleted of SNVs (Figure 3B). This heterogeneity extended to other measures of variation, such as nucleotide diversity (*π*). Genome-wide nucleotide diversity likewise varied by *>*100-fold among populations and substantially within and among genera (Figure 3C). In line with the ecological observations made above, we did not observe any significant differences in measures of genetic variation across discrete seasonal transitions. Despite these large intrinsic differences regarding genetic variation within microbial populations, we wondered how this variation changed over the course of our sampling. To measure how subspecies variation was partitioned over time, we calculated genome-wide temporal *F*_*ST*_. *F*_*ST*_ differed substantially within and among genera, with individual populations exceeding values of 0.4 (Figure 3D). With our observations above, we find that persistent bee-associated bacterial populations differ dramatically in both standing diversity and the extent of genetic differentiation through time.

**Figure 3.**
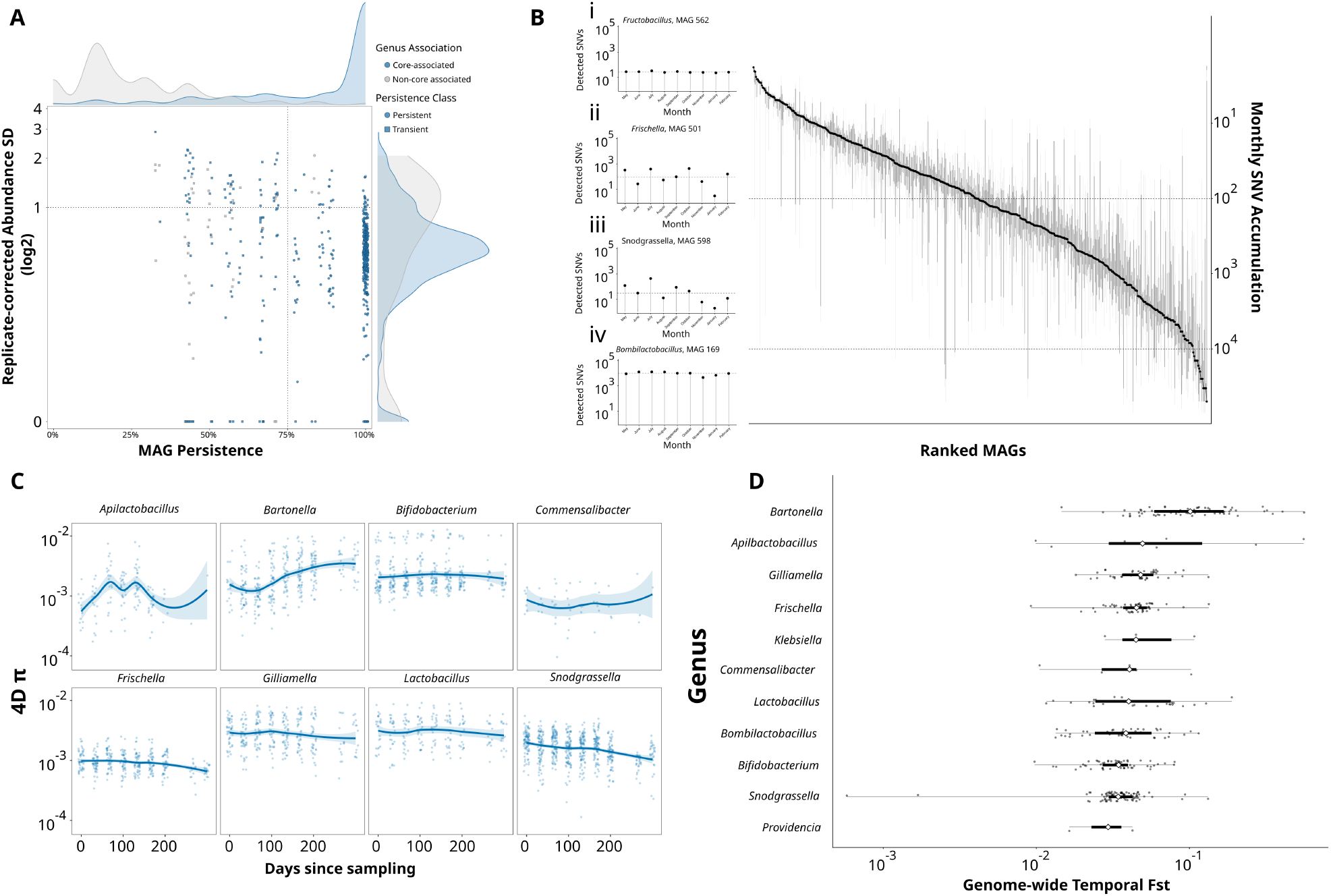
**A)** Persistence and replicate-corrected variance in temporal abundance for microbial populations exhibiting high coverage. Populations that belong to previously established core genera are colored blue; all others in gray. The relative density of both population persistence and temporal abundance are shown by the density distributions on either axis. **B)** Monthly SNV accumulation (y-axis) by population (x-axis). Black points denote median SNV accumulation across all sampled timepoints while the gray bars demarcate the interquartile range (central 50% of range) for all recovered microbial populations. Insets describe the number of detected SNVs across each month for a subset of individual microbial populations. The dotted line within each inset describes the median number of detected SNVs. **C)** Synonymous nucleotide diversity (y-axis) throughout sampling for select genera. Individual dots represent population-level averages. **D)** Genome-wide temporal *F*_*ST*_ (x-axis) for microbial genera with at least 10 representative populations (x-axis). White diamonds represent the median *F*_*ST*_ for each genus while individual gray points represent microbial populations.

### 3.4 Standing genetic variation is constrained by strong purifying selection

To better understand the contribution of natural selection on the observed standing genetic variation within bee-associated microbiomes, we quantified nonsynonymous and synonymous polymorphism within each MAG x colony x timepoint combination. Treating *π*_*S*_ as a proxy for neutral diversity and genealogical depth (see Methods and [17], *π*_*N*_ */π*_*S*_ declined strongly with increasing *π*_*S*_ (Figure 4A) with genetically diverse populations exhibiting substantially lower relative nonsynonymous diversity. This relationship was robust across multiple models (Extended Data 6; see Methods), including a modified model of time-dependent purifying selection (see [17]) in which deleterious nonsynonymous variants are removed at a rate relative to their fitness cost.

**Figure 4.**
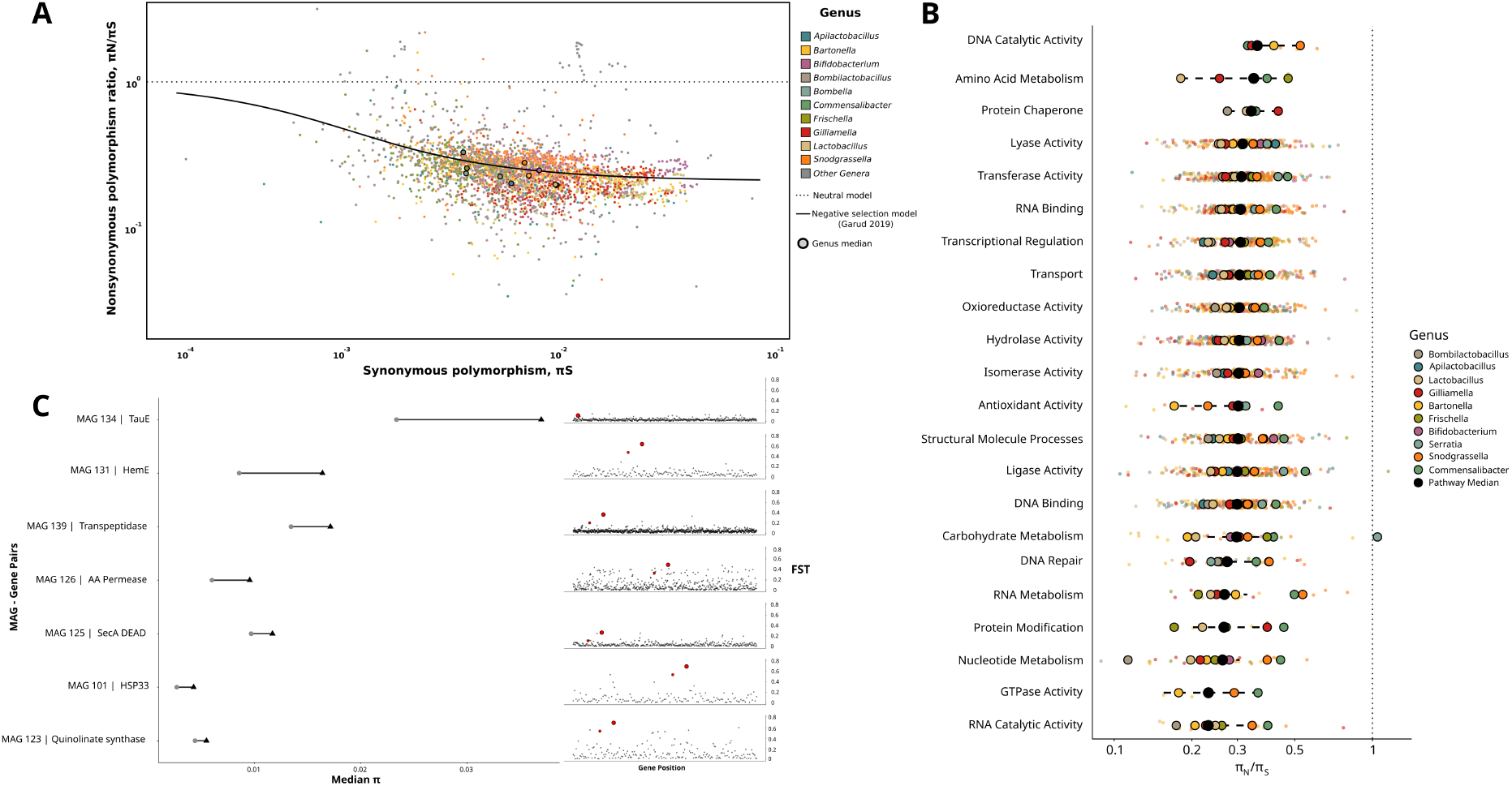
**A)** Ratio of diversity at nonsynonymous sites (*π*_*N*_) and synonymous sites (*π*_*S*_) as a function of *π*_*S*_ for all MAGs *×* colony *×* timepoint. Highlighted circles are representative of genus-level medians. The dotted line represents a theoretical prediction based on neutral processes only while the solid black line represents a prediction from a modified purifying selection null model seen in [17]. **B)** Mean *π*_*N*_ */π*_*S*_ values for different biological pathways (y-axis). Smaller colored dots represent MAG-specific pathway means while larger, highlighted dots represent genus-specific means. The dotted line centered at 1 represents a theoretical prediction based on neutral processes only. Black dotted lines within each pathway show bootstrapped 95% confidence intervals. **C)** Examples demonstrating genes enriched for *F*_*st*_ and nucleotide diversity (*π*). For each MAG *×* gene pair (y-axis, left figure), the median *π* value is shown (black triangle) compared to the genome-wide median (gray circle). The corresponding *F*_*st*_ value for this gene is shown (right figure) as a red circle relative to other genes in the MAG.

We next asked whether genes associated with specific pathways deviated systematically from this population-specific *π*_*N*_ */π*_*S*_ baseline. As expected, pathways associated with essential cellular processes exhibited comparatively low *π*_*N*_ */π*_*S*_, whereas several pathways associated with metabolism, lyase activity, and transport routinely exhibited *π*_*N*_ */π*_*S*_ values greater than expected (Figure 4B). Despite this heterogeneity, pathway-wide medians generally remained below the neutral expectation, emphasizing that functional differences occur against a genome-wide background dominated by purifying selection. Deviations were heterogeneously distributed among bacterial genera, which we explored further through measures of temporal genic *F*_*ST*_ (Figures 4C). Overall, we observed that genes associated with higher *F*_*ST*_ were more frequently associated with pathways exhibiting greater relaxed selection relative to expectation. Our results suggest that nonsynonymous variation in bee-associated microbes can be explained by strong purifying selection and that a subset of functions routinely deviates from this constrained background.

### 3.5 Rapid changes in allele frequency are frequent within bacterial populations

Against this background of strong purifying selection, we identified SNVs undergoing large changes in allele frequency between consecutive observations (e.g. *<* 20% to *>* 70%; see Methods). Because these events can result from selection, genetic drift, demography, or turnover among genetically distinct lineages, we refer to them as rapid allele-frequency shifts (Δ*RAF*). While distinguishing the relative contribution of these processes towards observed changes in allele frequency remains a significant methodological challenge, we sought to better understand how Δ*RAF* events were distributed along a continuum of localized genetic change to broader population turnover.

Individual populations associated with a range of Δ*RAF* dynamics (Figure 5A). For example, *Gillamella* MAG 254 exhibited localized Δ*RAF* with little abundance change, *Lactobacillus* MAG 191 exhibited broad concurrent shifts accompanied by an approximately 100-fold abundance change, and *Commensalibacter* MAG 411 showed repeated shifts against a comparatively retained genomic background (Figure 5A). Nearly all observed Δ*RAF* events were observed in replicate samples taken at each timepoint (Extended Data 7). Rather than falling into discrete categories of evolutionary or ecologically-mediated processes, these examples suggest that Δ*RAF* dynamics span a continuum from localized genetic modification to broad genomic change associated with large changes in population abundance.

**Figure 5.**
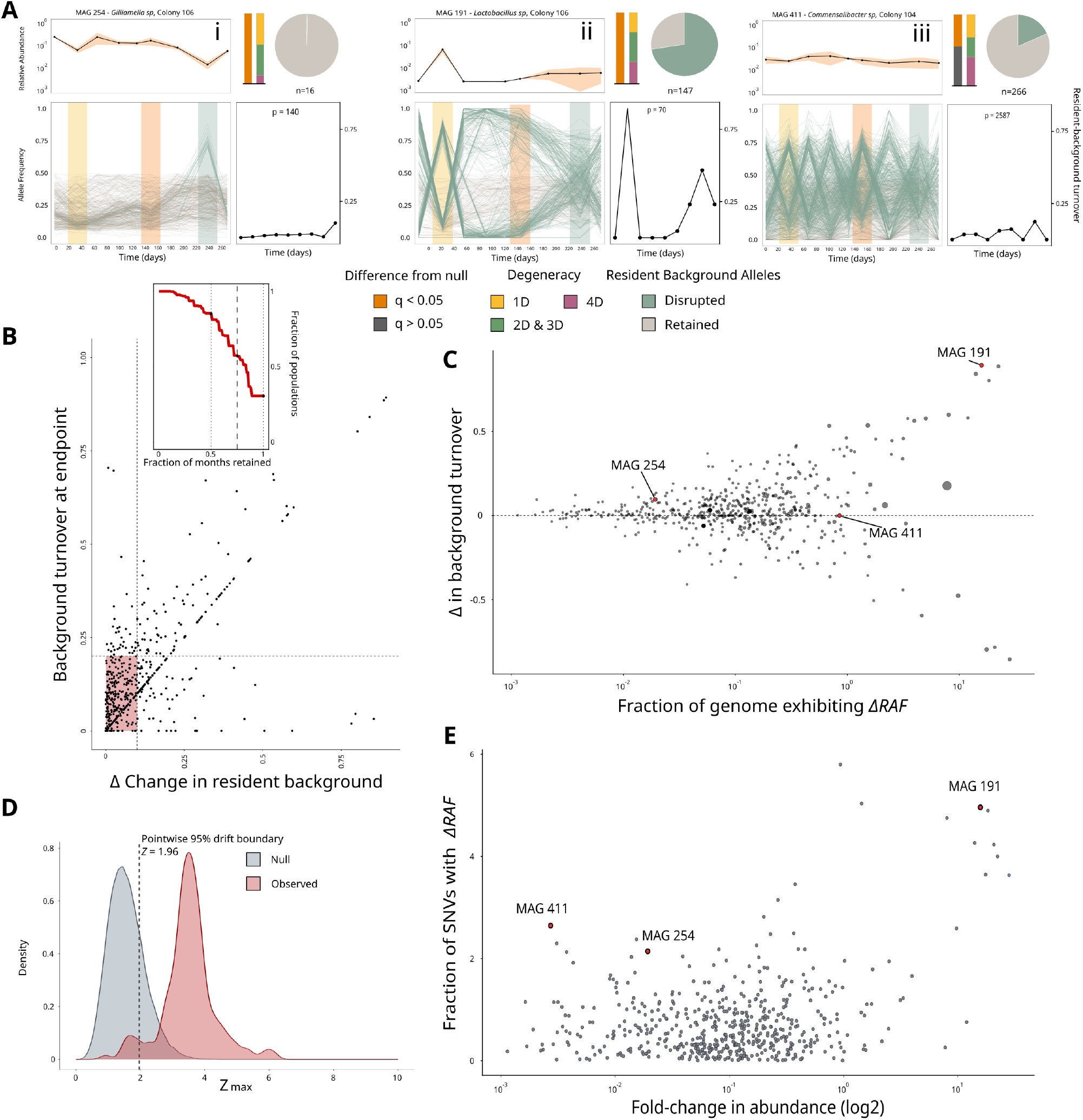
**A)** Representative patterns of rapid changes in allele frequency across three example MAGs. Upper subpanels: MAG relative abundance within its colony over time. Lower subpanels: Individual allele trajectories, highlighting rapid frequency shifts (blue) versus background alleles (subsampled gray). Seasonal transitions are indicated by yellow, orange, and blue background shading. Circular insets show the proportion of population SNVs exhibiting large frequency shifts. Adjacent bar plots show the proportion of SNVs exceeding a genetic drift null model (left, gray/orange) and their predicted codon degeneracy (right). Line plots indicate the proportion of near-fixed *T*_0_ alleles exceeding the frequency-shift threshold. **B)** Shift in resident genomic background during a focal allele sweep (x-axis) relative to total background shift at the final timepoint (y-axis). Pink shading highlights populations undergoing focal sweeps without substantial background turnover. Inset: Fraction of observed months (x-axis) versus the proportion of populations retaining their initial background (y-axis); dashed line marks 75% retention. **C)** Proportion of population-specific SNVs with rapid frequency shifts (x-axis) versus resident background turnover (y-axis). Circle size reflects the count of rapidly shifting alleles; red circles denote example MAGs from panel A. **D)** Abundance fold-change (log_2_-scaled, y-axis) relative to the fraction of SNVs exhibiting significant allele frequency shifts (x-axis). **E)** Standardized distribution of observed allele frequency shifts (pink) relative to a genetic drift null model (gray).

To quantify this continuum, we tracked near-fixation marker alleles from the initial population observation to calculate a background-turnover score (*B*_*t*_), where *B*_*t*_ *≈* 0 is indicative of retention of the initial background. Across 478 populations, the median population retained its initial background during 83.3% of sampled months (Figure 5B, inset). Crucially, 60.1% of Δ*RAF* intervals were resident-preserving (*B*_*t*_ *≤* 0.20, Δ*B*_*t*_ *≤* 0.1, cluster bootstrap 95% CI: 54.7-65.3%), and 73.4% of populations with Δ*RAF* contained at least one resident-preserving event.

Although Δ*RAF* events sometimes coincided with the disruption of dominant population-specific genetic backgrounds (e.g. strain haplotypes), we observed that most of this genetic variation occurred within highly stable demographic processes. First, we tested whether we could associate large numbers of simultaneous Δ*RAF* events with disruptions of the genetic background. The genome-wide extent of Δ*RAF* event – measured as the fraction of trackable SNVs undergoing rapid change – was positively correlated with the turnover of the initial resident background, Δ*B*| (*ρ* = 0.449, cluster bootstrap 95% CI, 0.370-0.532, Figure 5C). In contrast, these genetic changes were largely uncoupled from fluctuations on population abundance *ρ* = 0.121, cluster bootstrap 95% CI, 0.024-0.226, Figure 5D). In fact, we observed that approximately 75.1% of Δ*RAF* events involved less than a twofold change in population-level abundance, and nearly half (49.1%) occurred while both population abundance and the dominant genetic background remained stable. Altogether, these findings support that substantial allelic sweeps frequently occur without conspicuous shifts in population abundance or large changes in the dominant genetic background.

To test whether drift could explain Δ*RAF* dynamics, we parameterized a conservative neutral model maximizing variation attributable to drift and sampling noise (see Methods and [17, 19]). Standardized trajectories (Z max) were strongly displaced from neutral expectations, with most exceeding | *Z* | = 1.96 boundary (Figure 5C and Extended Data 8). While rejecting neutrality does not prove adaptation, drift and sampling uncertainty alone cannot account for the majority of Δ*RAF* events.

### 3.6 Recurrent nonsynonymous changes in allele frequency highlight candidates for adaptive evolution

Having identified a large subset of Δ*RAF* that were not readily attributable to common ecological or neutral explanations, we sought additional signatures that would independently strengthen the evidence for adaptive evolution. We focused on Δ*RAF* occurring at 0-fold degenerate sites (0*D* Δ*RAF*), where any allelic variation results in an amino acid change. We then asked whether these events were associated with local deviations in genomic diversity akin to what is observed in selective sweeps.

For each focal 0*D* Δ*RAF*, we identified all trackable synonymous SNVs (4-fold degenerate sites or 4D) across different genes located on the same contig and measured their absolute change in allele frequency during the focal event. Synonymous change was expressed relative to the median 4D change within the corresponding *MAG x Colony x Timepoint*) to account for differences in genome-wide dynamics in allele frequency change (see Methods). Excess synonymous change was greatest immediately surrounding focal 0*D* Δ*RAF* events and declined with increasing distance from the 0*D* Δ*RAF* site, approaching the genomic background over approximately 10-25 kb (Figure 6A). Similar short range covariance was also detected around nonsynonymous loci that did not experience a rapid shift in frequency, indicating that spatial covariance was not an artifact specific to 0*D* Δ*RAF* events.

**Figure 6.**
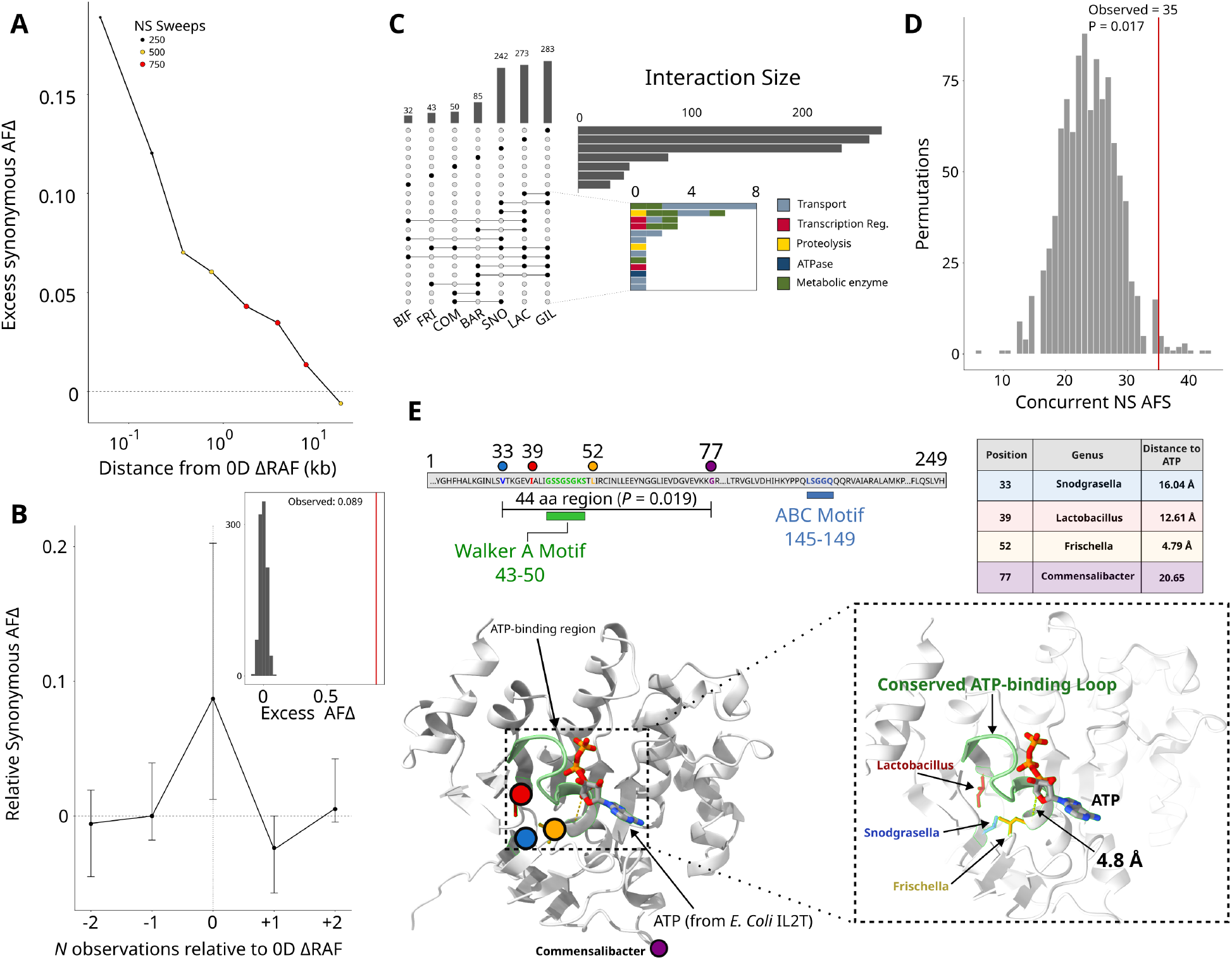
**A)** Lineplot showing the excess change in synonymous allele frequency (y-axis) relative to distance from a focal rapid change in allele frequency at a nonsynonymous site (x-axis). The dotted line centered at *∼* 0 shows the expected change at synonymous sites. **B)** Observations of neighboring synonymous allele frequency change (y-axis) at timepoints prior, concurrent, and following a focal change in NS allele frequency (x-axis). The timing of the focal event is centered at *x* = 0. The inset shows a distribution built by permuting the median expected allele frequency across rapidly changing sites compared to the observed change during changes in NS allele frequency (red line). **C)**.Upset plot showing the number of rapid changes in NS allele frequency per genus. Intersections among multiple genera represent examples where a rapid change occurred in genes found within a homologous protein cluster contained in multiple populations and occurring within the same month and across different colonies. The inset above contains a histogram showing how these functions are partitioned across different functional groups. **D)** Distribution built by permuting the observed number of 0*D* Δ*RAF* loci independently within each MAG x colony x time interval, normalized by the opportunity within for 0*D* Δ*RAF* events to occur across genes. The observed number of events is shown at the red line. **E)** Four independently observed 0*D* Δ*RAF* events from distinct microbial genera were identified via alignment of a representative ABC Transporter ATP-binding protein. The respective Walker A motif and ABC motif are highlighted in green and blue, respectively. The adjacent table shows the relative residue position, Genus, and distance to predicted ATP-associated phosphate for the identified variants. Colors denote independent microbial populations. Residue identities shown on the representative structure correspond to the medoid sequence rather than the population-specific ancestral or derived amino-acid states.

However, spatial covariance alone is unable to distinguish interval-specific rapid change from other genomic regions that are consistently variable through time. We therefore asked whether the same local regions became unusually dynamic specifically during timepoints containing focal 0*D* Δ*RAF* events. For each event, we followed nearby synonymous variants within 5 kb across all available timepoints and compared their median change in allele frequency with the typical temporal behavior of the same region. Local synonymous change increased sharply during the focal interval (*t* = 0) while preceding and subsequent intervals remained near the region-specific baseline (Figure 6B). The observed median excess in local synonymous change was 0.089, compared to approximately zero under the permutation null (*P* = 0.001). Further, we observed that the temporal interval associated with 0*D* Δ*RAF* exhibited greater allele frequency change in approximately 50% of events, compared with approximately 15% under the null (Figure 6B, inset). These findings suggest that increased variation in regions harboring 0*D* Δ*RAF* sites is not due to some innate bias towards variation within the region, rather that they exhibit rapid variation during the same time interval as the focal 0*D* Δ*RAF* event.

Because linkage-based signatures remain difficult to interpret without resolved haplotypes, we next asked whether 0*D* Δ*RAF* independently recurred within homologous proteins across microbial populations. Retained protein sequences were clustered into sequence-defined families (see Methods), revealing recurrent 0*D* Δ*RAF* across multiple MAGs, genera, and colonies (Figure 6C). We defined the strongest events as temporally concurrent when homologous proteins experienced 0*D* Δ*RAF* during the same exact sampling interval in at least two MAGs, genera, and colonies.

An opportunity-weighted permutation preserving the observed 0*D* Δ*RAF* counts within each MAG × colony × interval yielded a null median of 24 concurrent events, compared with 35 observed (*P* = 0.017; Figure 6D). Thus, only a small fraction of the much larger ΔRAF set satisfied our stringent criterion for independent temporal recurrence across microbial populations. This finding suggests that populations from different bacterial genera repeatedly experience nonsynonymous change in homologous proteins during the same sampling interval more often than expected from mutational opportunity alone. As a final analysis, we examined the taxonomic and functional distribution of the strongest recurrent protein families. Candidate protein families were distributed across multiple bacterial populations and included homologs related to transport, metabolism, protein homeostasis and transcriptional regulation (Extended Data 9). Several families experienced 0*D* Δ*RAF* events across multiple genera and colonies during the same sampling interval, which suggests that taxa exhibiting broad phylogenetic distance exhibit functionally similar targets of adaptive processes. For example, in one set of ABC transporter homologs, we observed four independent 0*D* Δ*RAF* across 4 genera mapped within a 44-amino acid region (*P = 0*.*019* vs permutation null). Three affected residues clustered spatially within a 13.2 °A on the predicted structure around a conserved ATP-binding loop, including an L*→* R substitution 2.9 °Afrom the Walker A motif and 4.8 °A from the ATP-associated phosphate (Figure 6E). Notably, three of the four affected residues were buried or nearly buried in the predicted structure, while the fourth was solvent exposed, suggesting that recurrent changes span both the structural core surrounding the nucleotide-binding region and a more accessible protein surface. Our observations of independent rapid nonsynonymous frequency shifts repeatedly localized to the same region of a conserved protein suggest that phylogenetically disparate taxa within the bee worker microbiome exhibit highly convergent responses to environmental change. Together, the spatial, temporal, and recurrence analyses narrow the much larger collection of rapid allele-frequency shifts to a restricted subset that we consider the strongest candidates for rapid adaptation within the honey bee gut microbiome.

## 4 Discussion

There is a growing consensus that microbiome genetic variation can change via adaptive processes within hosts over rapid and ecologically relevant timescales [17–21, 24, 44–47]. Most studies of host-associated microbial evolution, however, have focused on humans or mouse systems. The comparatively simple honey bee microbiome provides a complementary system in which genome-resolved longitudinal metagenomics allowed us to simultaneously track ecological and genetic change across hundreds of bacterial populations. We found that persistent populations harbored extensive standing genetic variation shaped predominantly by strong purifying selection, while a subset of nonsynonymous alleles underwent rapid, localized, and recurrent changes. At the community level, our findings are consistent with previously described differences between summer and winter microbiomes [38, 39, 48, 49], but suggest that community composition changes primarily across continuous time rather than through discrete seasonal states. This turnover was partially synchronized among colonies and more strongly synchronized within apiaries, yet concentrated among relatively few bacterial populations.

The distinction between ecological and evolutionary change becomes especially blurred below the level of microbial species, where lineage turnover, recombination, drift, and selection can generate similar allele-frequency trajectories. We therefore treated rapid allele-frequency shifts as a continuum rather than assigning each event to discrete “ecological” or “evolutionary” categories. Broad genetic changes increasingly coincided with resident-background turnover, yet many rapid shifts occurred while both population abundance and the resident background remained comparatively stable and exceeded conservative expectations from drift and sampling uncertainty. Our strongest evidence therefore comes from combining qualitatively independent signatures. Rapid nonsynonymous shifts were associated with spatially and temporally localized changes at neighboring synonymous variants, while homologous protein families experienced temporally concurrent nonsynonymous change across distinct microbial populations more often than expected from mutational opportunity. The recurrence of similar protein targets across phylogenetically distinct community members raises the possibility that multiple bacterial populations experience shared selective pressures within the host environment. As an example, transport systems repeatedly appeared in our analyses as common targets of putatively adaptive processes. The prevalence of ABC transporters, especially, highlights their role across disparate bacterial taxa in responding to shifts in microbial environments or host physiology, caused by changes in host diet or seasonal variation in plant-encoded secondary metabolites. Further studies seeking to explore how bee-associated microbes respond to changes in host diet or physiology, might consider interrogating these broad class of proteins. Together, these analyses narrow a much larger collection of rapid genetic changes to a restricted subset with multiple signatures consistent with adaptive evolution.

Several limitations arise from the stringent filtering required to minimize errors intrinsic to complex metagenomic data. Our pipeline excluded rare or unevenly covered MAGs, variable-copy genes, and highly similar genes shared across assemblies, likely removing some biologically important loci. Our focus on large frequency changes also misses gradual responses, soft sweeps, and variants experiencing weaker selection or clonal interference, while short-read sequencing prevents direct resolution of haplotypes. Our sampling did not capture a complete annual cycle, further limiting inference about recurring seasonal effects. Sample-matched isolates and long-read sequencing will therefore be important for resolving co-occurring lineages and directly testing the fitness consequences of candidate variants. Despite these limitations, our results demonstrate that persistent microbiome populations can undergo substantial evolutionary change on ecologically relevant timescales and that combining ecological context, resident-background stability, conservative neutral expectations, and independent recurrence provides a tractable framework for prioritizing candidate adaptation in natural microbial communities.

## 5 Methods

### 5.1 Sampling, Sample Processing, and Sequencing

Twelve untreated honey bee colonies across three apiaries in Illinois and Indiana were sampled monthly from May 2023 to March 2024. From each colony, in-hive bees (IHBs; *n* = 60 dissected across 3 pools of 20) and late-instar larvae (L4/L5; 5 individual and 1 pooled sample) were collected. IHB gut tissue was homogenized and fractionated by differential centrifugation to enrich the microbial fraction [50]. Total gDNA was extracted using Qiagen DNeasy UltraClean Microbial kits, libraries prepared with BioNEXTflex Rapid DNA Library kits, and paired-end sequencing performed on an Illumina NovaSeq X. Detailed field sampling and laboratory protocols are available in Supplemental Text S1.

### 5.2 Metagenomic Assembly, Binning, and Taxonomy

Quality filtering was performed using bbduk [51]. Filtered reads were co-assembled across replicate samples using MEGAHIT [52] and mapped back using bowtie2 [53] to generate BAM files [54]. Metagenomic bins were reconstructed using VAMB [55] and MetaBAT2 [56], integrated via Binette [57], and evaluated with CheckM2 [58]. Bins meeting quality thresholds (*>* 50% completeness, *<* 10% contamination; *n* = 1,028) were taxonomically classified with GTDB-TK [59] and dereplicated into reference MAGs (rMAGs) at 95% ANI using dREP [60, 61]. Full command-line parameters are provided in Supplemental Text S1.

### 5.3 Colony-Specific Mapping Databases

To consistently quantify allele frequencies and clade abundance through time while preventing ambiguous read partitioning among closely related lineages, we constructed colony-specific reference databases rather than mapping reads to one large database [17, 62]. An rMAG was included in a colony’s database if and only if it contained a dereplicated MAG originally recovered from that focal colony.

### 5.4 Metagenomic pipeline for ecological analyses

#### 5.4.1 rMAG Quality Control, Abundance, and Persistence

To prevent assembly artifacts and off-target read mapping from heavily biasing downstream ecological metrics, reconstructed reference MAGs (rMAGs) were filtered based on genome size (*<* 10 MB), contig recrutiment breadth (*≥* 25% of contigs recruiting reads in a sample), and coverage depth (median coverage *≥* 5 or mean and median coverage both *>* 1). Reads were mapped to colony-specific databases to estimate average genome-wide coverage and to calculate sample-level relative abundances. To assess longitudinal occupancy within a colony (*O*_*ic*_), an rMAG was scored as detected in a colony-month (timepoint or interval) if it satisfied quality criteria in *≥* 50% of biological replicates. Populations observed across *≥* 4 eligible months were classified as persistent if detected in *≥* 75% of eligible months (including the final two months). Populations were otherwise classified as intermittent. Replciate-corrected temporal variance (*σ*^2^) was calculated by partitioning within-timepoint replicate variance from variation in longitudinal abundance.

#### 5.4.2 Ecological Diversity Analyses

Alpha diversity metrics were calculated using *vegan* [63]. Group differences were evaluated via Kurskal-Wallis tests and Dunn’s post-hoc tests utilizing Benjamini-Hochberg (BH) correction for false discovery rate. Differences in sample size among seasons were controlled for via subsampling (B = 5000, N = smallest seasonal sample size). For cross-colony beta diversity analyses, rMAG abundances were first aggregated to genus-level abundances, square-root transformed, and analyzing using Bray-Curtis dissimilarity. Distance-based redundancy analysis (dbRDA) conditioned on colony identity (Condition(colony)) was performed alongisde PERMANOVA, restricting permutations within colonies to account for repeated observations due to longitudinal sampling. To isolate seasonal transitions from cumulative time lag, residual dissimilarity (*R*_*ij*_) relative to temporal distance was evaluated with colony-level circular shift permutation.

#### 5.4.3 Community Turnover, Apiary Synchrony, and Taxon Enrichment

Consecutive-month turnover (*D*_*c,t*_) was calculated as the Bray-Curtis dissimilarity between consecutively sampled months within each colony. Inter-colony synchrony scores were evalulated using pairwise Spearman correlations (*r*_*ij*_) of *D*_*c,t*_ trajectories and comparing of intra-apiary versus inter-apiary coordination against a null distribution (B = 5000). Total colony turnover was then decomposed into individual MAG fractional contributions (*C*_*i*_), and the minimum proportion of active MAGs explaining 50% (*N*_5_0) and 80% (*N*_8_0) of total turnover was extracted from cumulative Lorenz curves. Genera disproportionally represented among dominant turnover contributors (*≥* 50% cumulative turnover) were identified using within-colony genus-label permutation tests with BH FDR correction. Full mathematical formulations and permutation mechanics, as well as QC metrics are detailed in Supplemental Text S!.

#### 5.4.4 Core Genome Definitions, SNV Profiling, and Population Diversity

Core genomes for each rMAG were defined by retaining DRAM-predicted genes [64] with consistent relative copy numbers (0.3 *≤ C*_*i,t*_ *≤* 3, *<* 10% sample failure rate) and excluding non-unique sequences exhibiting *≥* 95% ANI across distinct rMAGs via MMseqs2 [17, 19, 65, 66]. Single nucleotide variants (SNVs) were profiled using Anvi’o [67] on rMAGs containing *≥* 100 core genes. SNVs were restricted to coding regions meeting coverage thresholds 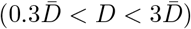 present in *≥* 75% of colony samples. Nucleotide diversity (*π*), Watterson’s *θ*_*w*_, Tajima’s *D*, and non-synonymous to synonymous site ratios (*pN/pS*) were computed across polarized 0-fold (0D) and 4-fold (4D) degenerate sites [41, 68, 69].

#### 5.4.5 Allele Frequency Dynamics, Genomic Backgrounds, and Divergence

Rapid allele frequency shifts (Δ*RAF* ; transitions from *<* 20% to *>* 70% frequency between consecutive timepoints) were identified using a binomial sampling error model (*FDR <* 0.1, *q <* 0.05) [17, 19] and validated against independent biological replicates. Temporal genetic divergence was quantified via genic *F*_*st*_ [70]. To distinguish *de novo* mutations from episodes of strain replacement, resident genomic backgrounds were tracked using high-frequency baseline marker alleles (*f*_resident_ *>* 0.9 at *t*_0_), defining displacement as *B*_*t*_ = 1 *−* median(*f*_resident,*t*_) [19]. Trajectories were classified as resident-preserving when *B*_*t*_ *≤* 0.20 and | Δ*B* |*≤* 0.10. Shift breadth (*S*) was evaluated as the proportion of trackable SNVs undergoing Δ*RAF* relative to population abundance changes.

#### 5.4.6 Selection Null Models, Parallelism, and Neutral Drift Models

To evaluate purifying selection without artefactual correlations due to shared denominators, the relationship between *π*_*N*_ */π*_*S*_ and *π*_*S*_ was modeled using a gene-splitting decoupling procedure [17, 66, 71]. Spatial and temporal covariance of 4D SNVs surrounding focal 0D Δ*RAF* sites (*≤* 5 kb) were evaluated against matched control loci and within-region permutations. To evaluate allele frequency shifts against neutral drift, minimal effective population sizes (*N*_*e*,min_) were estimated per MAG from 4D variance under Wright-Fisher diffusion [19], yielding standardized score trajectories 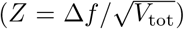 and trajectory maxima (*Z*_max_).Parallel evolution was assessed by clustering core proteins into families with MMseqs2 (50% minimum identity, 85% coverage) [65]. Recurrent 0D Δ*RAF* events across distinct MAGs and colonies were tested against a gene-length-weighted permutation null (*B* = 5, 000) [18, 20, 21], with seasonal rates modeled with generalized linear mixed models. To evaluate within-protein structural localization, members of recurrent candidate families were aligned with MAFFT, and original 0D Δ*RAF* positions were mapped onto representative medoid sequences. Significance for spatial clustering was tested for via random permutation of residue labels across populations. For a focal ABC Transporter ATP-binding protein, the 249-aa medoid sequence was structurally predicted using AlphaFold2-ptm [72] (ColabFold v1.6.2 [73]). Pairwise C*α* and minimum heavy-atom distances were calculated from the predicted coordinates to confirm the structural clustering of independently observed events (e.g., positions 33, 39, and 52 forming a compact region spanning 13.24 °A). Finally, minimum heavy-atom distances were calculated between these mapped positions, the conserved Walker A motif, and an ATP ligand, the latter modeled by superimposing the medoid structure onto the experimentally determined ATP-bound dimer 1L2T using UCSF ChimeraX [74]. Detailed mathematical formulations, parameterizations, and structural mappings are provided in Supplemental Text S1.

## Acknowledgments

We thank Nathan Beach for his helped in maintaining colonies at UIUC during the sampling period. We are grateful to A. Murat Eren and the Helmholtz Institute for Functional Marine Biodiversity for providing work space and feedback during the building of the analytical pipeline. We are grateful to David Merrit and the Center for Genomics and Bioinformatics at Indiana University for their help with gDNA extraction and library preparation. LLMs were used to generate code and build out a portion of the bioinformatic pipeline. This work was financially supported by a scholarship from the International Symbiosis Society and a Costco/Project Apis m. research grant to C.R.P.R., an NSF IOS Collaborative Research award (2005306) and by an NSF DBI Biology Integration Institutes award (2022049) to I.L.G.N.

## Data availability

Raw metagenomic reads are available from NCBI under PRJNA1531180. Raw code, scripts, and workflows are available here: https://github.com/en-nui/BEE-WITCH

