## SupplementalMethods for "Ecological stability masks rapid, convergent evolution within the honey bee worker microbiome"

**This PDF file includes:**

Supplementary Methods

Supplemental Figures and legends S1 to S20

Legends for Supplemental Tables S1 to S2

### Supplementary Methods

#### 0.1 Study design and sampling regime

12 honey bee colonies were maintained across three apiaries in Illinois and Indiana. Colonies were maintained by professional beekeeping staff but were otherwise untreated for common hive parasites (e.g. *Varroa* and small hive beetle). Colonies were sampled monthly from May 2023 to March 2024 or until colony death. Sampling was conducted as follows: frames near the center of the hive were temporarily removed and checked for the presence of the queen. In-hive bees (IHB) were then transferred to large plastic containers via vigorous shaking of the frame until approximately 100 IHBs were recovered. All IHBs at time of sampling came from a single frame. Two colonies were further sampled for larvae. For larval sampling, we sampled IHBs as before, but ensured that the IHBs were recovered from a brood frame. Late instar larvae (L4 and L5) were removed from cells via forceps and placed into individual Eppendorf tubes until 5 individual larvae were collected. One final tube was collected consisting of 5 pooled larvae. IHBs and larval samples were stored on ice until processing at the University of Illinois Urbana-Champaign or Indiana University.

#### 0.2 Sample processing, DNA Extraction, Library Prep, Sequencing

IHBs were placed at 4C until incapacitated. Gut tissue from individual IHBs was dissected and placed into a single Eppendorf tube until 20 tissue samples had been collected. This process was repeated two more times, resulting in the dissection of 60 IHBs split across 3 pools of 20 individual IHBs. Following dissection, pooled tissue was transferred to a sterile Dounce homogenizer and homogenized on ice, as performed in 1. 3 ml of homogenate was pipetted into 3 sterile 1.5 ml microcentrifuge tubes and spun at 500G for 10 minutes to pellet host tissue. Supernatant from all 3 tubes was strained through a 40 $\mu M$ cell strainer to remove the majority of host cells and collected in a sterile Falcon tube. 1.5 ml of the pooled supernatant was then transferred to a new microcentrifuge tube and spun down at 5000G for 10 minutes to pellet the microbial fraction. The supernatant was then removed and the remaining pellet was flash frozen and stored at -80C. Larval samples were homogenized with a plastic pestle before being flash frozen and stored at -80C. Samples were processed for gDNA extraction and library prep at the Center for Genomics and Bioinformatics at Indiana University. gDNA extraction was carried out with Qiagen DNeasy UltraClean Microbial kits and libraries prepared with BioNEXTflex Rapid DNA Library kits. DNA was sequencing as paired end libraries on a 25B Flow cell and ran on a NovaSeq X at the Indiana University School of Medicine's Center for Medical Genomics.

#### 0.3 Read QC, Assembly, mapping binning strategy and GTDBTK

Read QC was carried out with bbduk [2] (ktrim=r k=23 mink=11 hdist=1 minlen=50 tpe tbo qtrim=r trimq=10 ftm=5 ftl=10). Processed reads were used as input for co-assembly with MEGAHIT [3] (megahit –presets meta-sensitive). Co-assembly was chosen to maximize the recovery of rare taxa and was carried out by pooling reads from replicate samples. Reads were mapped to assembles with bowtie2 (bowtie2 –very-sensitive) [4] and converted to BAM files [5]. Coverage files and assemblies were used as input for our binning pipeline, which combined bins VAMB [6] and Metabat2 [7] before final processing with Binette [8]. CheckM2 [9] was used to assess bin quality, completeness, and contamination. 1,028 bins with completeness >50% and contamination <10% were retained. GTDB-TK [10] was used to assign taxonomy to each bin (gtdbtk classify_wf). These bins were dereplicated at 95% ANI using dREP [11] with primary clustering at 80% and secondary clustering at 95% and 85% breadth. We ran dREP from 90% to 99% and examined the number of dereplicated bins as well as the number of genomes contained in each dereplicated bin. We chose 95% ANI because this maximized both the number of recovered bins, maintained the highest mean dREP cluster size, and has been repeatedly shown to generate species-level delineation for MAGs [12].

#### 0.4 Creation of rMAGs and custom mapping databases

To quantify abundance and allele frequencies consistently through time within colonies while reducing ambiguous recruitment among closely related genomes, we created custom genome databases for each colony in lieu of recruiting reads to all dereplicated MAGs. This was done in order to minimize reads being split among genes found on highly similar rMAGs, despite those rMAGs not being present in the focal colony. This approach has been applied in other studies to control for similar confounding effects [13, 14]. For each colony, we identified all MAGs that were assembled from that colony prior to dereplication. We then referenced this list of colony-specific MAGs to the recovered rMAGs during dereplication. As rMAGs often collapsed MAGs originating from multiple independent colonies, an rMAG was only included in a colony-specific database if it contained an MAG from that colony. Put another way, if an rMAG collapsed MAGs from both Colony A and Colony B, it would be present in the colony-specific database for both Colony A and Colony B. However, if an rMAG dereplicated an MAG from Colony A and not Colony B, it would only be present in the database for Colony A and not in the database for Colony B. This approach allowed us to partition genomes for read recruitment based on dereplication patterns that originated in the data, rather than relying on an all-vs-all approach or by arbitrary genome selection.

#### 0.5 Metagenomic pipeline for ecological analyses

##### 0.5.1 mMAG quality control for ecological analyses

Quantification of microbial abundance can be highly skewed by assembly artifacts, copy number variation, and off-target mapping. We sought to control for potential errors through the following steps:
1) rMAGs with predicted genome sizes $\geq$ 10,000,000 bp were flagged and removed. Chimeric assemblies are ubiquitous in metagenomic assemblies, and we sought only to remove the most egregious of cases as these are likely to represent multiple concatenated and highly chimeric genomes.
2) For each rMAG, we counted the number contigs associated with each assembly and compared it to the number of contigs that actually recruited reads after mapping to the colony-specific database. rMAGs were flagged in a sample if $<$ 25% of the total contigs associated with the rMAG were found to recruit reads.
3) rMAGs that passed steps 1 and 2 were then checked for coverage. rMAGs were kept when median coverage $\geq$5 or when both mean coverage and median coverage were $>$ 1. This step was implemented to control for MAGs where coverage counts were heavily skewed by genes common in the community (e.g. transposons) while the rest of the genome recruited little coverage.

rMAGs that failed the first filter (1) were removed completely from downstream analyses while those that failed filters 2 and 3 were typically removed due to the more stringent filters in downstream analyses (see below).

##### 0.5.2 Relative abundance estimates

To estimate the relative abundances of rMAGs in a given sample, reads from each sample were mapped to a colony-specific database dependent on the sample origin. Average genome-wide coverage was used to quantify the relative abundance of a given mMAG in a sample. This procedure was repeated for colony-wide estimates of each MAG, except that we normalized the coverage of each rMAG by the number of replicate samples in which it was present. These estimates were used for the relative abundance trajectories in Figures 2 and 3. Relative abundance was calculated independently within each replicate sample as:

$$P_{is} = \frac{A_{is}}{\sum_f A_{js}} \tag{1}$$

where $A_{is}$ is the abundance of MAG $i$ in sample $s$. Because MAG coverage was derived from colony-specific reference sets and were therefore not treated as a common feature among colonies, MAG relative abundances were summed by genus to generate a shared sample x genus matrix for analyses that utilize cross-colony ordination and beta-diversity analyses (see below). Biological replicate pools were retained as separate observations for sample-level ordination and were used for analyses explicitly comparing intra-colony dynamics.

##### 0.5.3 Longitudinal MAG detection and persistence

Persistence analyses were performed for each MAG x colony combination using thresholds specific to each colony. MAGs were retained if they met the presence/absence criteria defined above (see mMAG quality control for ecological analyses) in at least one valid worker metagenome. For each retained MAG associated with some colony, we quieried whether this MAG appeared in all eligible metagenomic associated with that colony. Missing MAG x library combinations were scored as non-detections. Nondetections were first evaluated inependently within biological replicate pools and then summarized within each colony-month combination. A MAG x colony combination was considered detected within a colony-month when it satisfied the presence/absence criteria in at least 50% eligible biological replicates. We then defined the longitudinal occupancy of some MAG population $i$ in colony $c$ as:

$$O_{ic} = \frac{number\ of\ detected\ colony - month\ combinations}{number\ of\ eligible\ colony - month\ combinations} \tag{2}$$

MAG x colony combinations observed in at least four eligible months were classified as persistent when they were detected in $\geq 75\%$ of eligible months and were detected within the first two and last two eligible months associated with that sample. All other populations were classified as intermittent.

##### 0.5.4 Replicate-corrected variation in abundance

To distinguish temporal changes in rMAG abundance from variation among biological replicates, we decomposed the observed variance for each MAG x colony abundance trajectory. For rMAG $i$ in colony $c$, replicate-level relative abundances were first averaged within each sampled timepoint $t$, producing $\bar{x}_{ict}$. Total temporal variance, $\sigma^2_{total,ic}$ was calculated across these averaged observations. We then estimated replicate-corrected temporal variance as:

$$\sigma^2_{\text{bio},ic} = \max\left(0, \sigma^2_{\text{total},ic} - \frac{1}{T_{ic}} \sum_{t=1}^{T_{ic}} \frac{\sigma^2_{\text{rep},ict}}{n_{ict}}\right) \tag{3}$$

where $\sigma^2_{\text{total},ic}$ is the variance among biological replicates at timepoint $t$, $n_{ict}$ is the number of replicate pools, and $T_{ic}$ is the number of eligible sampled timepoints. Negative estimates were truncated at zero. This statistic therefore represents temporal abundance variation after accounting for uncertainty among biological replicates.

##### 0.5.5 Estimates of community diversity

Richness, Simpson, and Shannon indices were generated for each sample, colony, season, and apiary using the R package vegan [15]. Differences in these metrics were compared using a Kruskal-Wallis test. In comparisons where this test was significant, pairwise comparisons were evaluated using Dunn's test with Benjamini-Hochberg false discovery rate (FDR) correction applied across all pairwise comparisons. To ensure that differences in sample size weren't a primary driver of the observed differences in alpha diversity among seasons, we sampled each season by $N = (\textit{number of samples in smallest season})$. During each sampling, we conducted a Kruskal-Wallis Rank Sum Test and constructed a distribution of the *p-values* with which we scored the percentage of significant iterations of the test. The robustness of a seasonal effect was defined as significant if >95% of p-values were less $< 0.05$. We then repeatedly sampled $N$ observations per season with replacement across 5000 iterations and calculated the mean Shannon, Simpson, and Richness measures for each season. These mean values were used to generate the respective distributions. For each iteration, we calculated $\Delta$, defined as the percentage of iterations where the mean of season $i$ is greater than the mean of another season:

$$\textit{Support} = \frac{\sum_{i=1}^{5000} I(\bar{x}_{A,i} > \bar{x}_{B,i})}{5000} \tag{4}$$

where $I$ is equal to 1 if the condition is true ($\bar{x}_{A,i} > \bar{x}_{B,i}$) and 0 otherwise. Values approaching 100% or 0% are indicative of strong support for seasonal differences among season. This approach allows us to address structural differences among data in the different seasons, which are likely to seriously affect the observed variance within seasons.

##### 0.5.6 Community ordination

As reads were recruited against colony-specific rMAG databases, individual rMAGs were not treated as a common feature for cross-colony comparisons. Corrected rMAG relative abundances were therefore summed by genus to generate a shared sample x genus abundance matrix. Biological replicates were retained as individual observations, and genus relative abundances were square-root transformed prior to calculating Bray-Curtis dissimilarity.

To visualize changes in community structure over time and indepently of colony-specific composition, we performed distance-based redundancy with *dbrda* in *vegan*, conditioning on colony identity:

$community \sim Condition(colony)$,

with Bray-Curtis dissimilarity as the response. The resulting constrained coordinates represent community variation after partialling out the component associated with colony identity. The proportion of total Bray-Curtis dissimilarity associated with colony metadata was quantified separately in a corresponding unconditioned model. Individual samples were colored according to continuous sampling time (e.g. month) for visualization. An unconditioned NMDS and PERMANOVA 15 analyzes of colony, apiary, month, season, and time lag were retained as supporting analyses. Temporal matadata were evaluated using permutations restricted within colonies to account for repeated longitudinal sampling. PERMANOVA $R^2$ values were interpreted primarily as descriptive effects as repeated sampling over time violates independence among samples.

##### 0.5.7 Bray-curtis dissimilarity over time

To quantify within-colony dissimilarity over time, genus-level relative abundances were first averaged across biological replicate pools from the same colony-month combination. Bray-Curtis dissimilarity was then calculated between all pairs of sampled months and belonging to the same colony. Temporal distance among pairs was calculated as the absolute difference in sampling time (as days) between the two observations. Each pair was described by its associated community dissimilarity $BC_{ij}$ and temporal distance $|t_i - t_j|$. Colony-level measurements of dissimilarity and time lag were for inference as replicate samples were non-independent.

##### 0.5.8 Comparing Bray-curtis dissimilarity over differences in time lag versus seasonal categories

Because seasonal progression is inherently correlated with time lag over the sampling period in our study, we tested whether differences in seasonality were associated with community divergence beyond what was expected from elapsed time alone. For each colony, Bray-Curtis dissimilarity was calculated at the genus-level among all colony-month combinations. The expected relationship between Bray-Curtis dissimilarity and time lag was estimated within each colony, and the residual dissimilarity was calculated as:

$$R_{ij} = BC_{ij} \; - \; \hat{BC}(|t_i \; - \; t_j). \tag{5}$$

Sample pairs were then classified as belonging to the same or different season. For each colony, we then calculated a statistic to capture the excess dissimilarity as a consequence of differences in season:

$$\Delta_{season,c} = \bar{R}_{different\ season} \; - \; \bar{R}_{same\ season} \tag{6}$$

where the overall statistic was the mean across colonies, thereby weighting colonies equally despite colony heterogeneity with respect to the number of observations.
Significance was assessed using circular shift permutation of seasonal labels within each colony. Circular shifting in this way preserves temporal ordering and the structure of each colony trajectory while disrupting the association between discrete seasonal categories and residual community dissimilarity. The observed statistic was compared with the resulting permutation distribution. Confidence intervals were estimated by resampling colonies with replacement.

##### 0.5.9 Magnitude of consecutive community change

To quantify the magnitude of ecological change during the time series, genus-level relative abundances were averaged across biological replicates within each colony-month and Bray-Curtis dissimilarity was calculated between consecutive sampling intervals (e.g. May to June, June to July, etc). The resulting statistic:

$$D_{c,t} = BC(p_{c,t}, p_{c,t+1}) \tag{7}$$

represents the magnitude of community change experienced by colony $c$ over time interval $t$.

##### 0.5.10 Synchrony of community turnover

To quantify whether microbial communities exhibited coordinated responses to seasonal change, we tested whether colonies tended to experience relatively large or small ecological changes during the same sampling interval. For each colony $c$ and consecutive sampling interval $t$, ecological turnover was defined as:

$$D_{c,t} = BC(p_{c,t}, p_{c,t+1}) \tag{8}$$

For each pair of colonies $i, j$, synchrony was defined as the Spearman correlation between $D_{c,t}$ magnitudes across sampling intervals shared by both colonies.

$$r_{ij} = \rho_{spearman}(D_{i,t}, D_{j,t}) \tag{9}$$

and apiary structure was summarized as:

$$\Delta = \bar{r}_{withinapiary} - \bar{r}_{betweenapiaries} \tag{10}$$

such that positive values indicate stronger synchronization among colonies housed within the same apiary.

To determine whether the observed apiary structure could arise from colony-specific temporal dynamics alone, we compared $\Delta_{apiary}$ under two simulated models. The null model preserved the temporal structure and variance associated with individual $D_{c,t}$ trajectories but omitted any apiary-shared component, while the fitted model retained apiary-associated covariance that was estimated from the observed data. For each permutation, $D_{c,t}$ trajectories were recalculated, converted to Colony x Colony Spearman-correlation matrices, and summarized using the same $\Delta_{apiary}$ statistic. This procedure was repeated $B = 5000$ times, and a one-sided p-value was computed as:

$$p_{\text{sync}} = \frac{\#\{S_{\text{null}} \geq S_{\text{obs}}\} + 1}{B + 1} \tag{11}$$

##### 0.5.11 Testing for contribution of individual MAGs to community turnover

Due to the observed positive correlation between community dissimilarity and increasing time lag, we tested whether individual MAGs had a disproportionate contribution to community turnover. Here, we define community turnover as the degree of intra-colony microbial community composition change across consecutive months. To decompose community turnover into contributions from individual MAGs, we used colony-level relative abundances for each MAG and calculated Bray-Curtis dissimilarity between two relative abundance values associated with any two consecutive-month pairs as:

$$BC_{t,t+1} = \frac{1}{2} \sum_{i} |p_{i,t+1} \ - \ p_{i,t}| \tag{12}$$

where $BC$ is Bray-curtis dissimilarity at timepoint $t$ and $p$ values reference the sample-level relative abundance at either timepoint. The contribution of MAG population $i$ to the total abundance turnover during sampling was defined as:

$$C_i = \frac{|p_{i,t+1} \ - \ p_{i,t}|}{\sum_j |p_{j,t+1} \ - \ p_{j,t}|} \tag{13}$$

MAGs detected at one or both endpoints of a consecutive-month pair were considered active contributors to community turnover. Populations absent at both endpoints were excluded from the denominator. Contributions sum to one for each monthly transition.

###### 0.5.12 Lorenz curves for community turnover

MAGs were ranked from largest to smallest values of $C_i$, and their cumulative contribution to community turnover was calculated for each consecutive-month sampling within each colony. Median and interquartile ranges were calculated across all month pairs. We calculated $N_{50}$ and $N_{80}$, defined as the smallest numbers of active MAG populations required to account for 50% and 80% of total turnover, respectively. These values were divided by the number of active populations within each month pair to derive the proportional breadth of community turnover.

###### 0.5.13 Testing for enriched taxa among dominant turnover contributors

For each consecutive month-pair within a colony, we defined the dominant turnover contributors as the smallest ranked set of active MAG populations whose cumulative abundance changes accounted for at least 50% of community turnover. For each MAG x colony population, we recorded the number of eligible month-pairs in which it was active as well as the number of pairs in which it was classified as a dominant turnover contributor. We then tested whether particular bacterial genera were disproportionately represented among dominant turnover contributors using a within-colony permutation test. Genus labels were permuted among MAG populations within each colony while each population's complete turnover trajectory was left intact. This null allows us to preserve the number of MAG populations within each colony, the genus composition of each colony, the number of month pairs associated with each population within each colony, and the contribution of each population to community turnover. For each permutation, the fraction of active MAGs within each consecutive month pair was classified as a dominant contributor was requantified for each genus. Observed genus-specific frequencies were then compared to their permuted distributions, and $p$-values underwent Bejamini-Hochberg correction.

#### 0.6 Metagenomic pipeline for evolutionary analyses

##### 0.6.1 Quantifying gene content within each MAG

To minimize the impact of sequencing errors and mismapping, we first quantified variation in coverage among genes in each MAG. We calculated the average and median coverage for each MAG in a given sample as well as for each individual gene predicted by DRAM [16]. Median coverage for each gene was calculated by using the gene coordinates predicted from the DRAM software. The copy number ratio of a given gene, i at some timepoint t, $C_{i,t}$, was calculated by dividing the median gene coverage by the genome-wide median coverage.

As in previous studies [14, 17, 18], we used the distribution of $C$ values to define a "core" genome for each MAG. Here, we define "core" by identifying genes that fit into two distinct classes. The first class pertains to genes that are stable within each MAG and exhibit similar coverage profiles relative to the genome-wide coverage of a MAG at some timepoint over the length of the sampling. To define this, we required genes to have coverage such that $0.3 \leq C \leq 3$ at a single timepoint. We used the total number of samples in a given colony that the gene appears to establish a baseline failure rate, defined as 10%. Genes that exhibited coverage deviations > 10% than the total number of samples in a given colony were flagged for removal.

The second class of genes are those that are likely not shared among different MAG_id, whether through HGT or misassembly. We used MMseqs2 [19] to reciprocally map genes from each rMAG in a colony. Genes that exhibited $\geq$ 95% ANI similarity between different rMAGs were flagged for removal. Genes that were included in both classes were used to define the "core" genome for each rMAG and used for all downstream analyses.

##### 0.6.2 Identifying SNVs within bacterial MAGs

Single nucleotide variant (SNVs) calling was based on a modified pipeline developed in [14, 18]. rMAGs that passed our ecological filters (see above) were used as input for Anvi'o [20]. Anvi'o-gen-variability pipeline was used to call SNVs in each MAG within each sample. By default, Anvi'o reports the coverage at each SNV and filters SNVs below 10x coverage. MAGs were excluded if < 100 genes were present in the core genome after gene filtration through the gene blacklist and gene whitelist. For all genes, we compared genic coordinates based on prediction and annotation from DRAM as well as predictions from Anvi'o. Any conflicts in coordinate overlap were removed, though this represented <1% of cases. Adopting methods from previous studies, we defined the total read coverage, $D$, for each site in a MAG. We then built coverage distributions across each MAG to obtain median $D$ for each SNV, $\bar{D}$.

MAG-specific estimates of  were used to refine our expectations for $D$ at any site in the genome. Sites with $D <<$  are likely to arise due to mapping errors or if the reference MAG contains genes that are absent in a given sample. Alternatively, SNVs with $D >>$  could be generated from genes of high copy number or read donation from other microbial species in a sample. To exclude either case, we kept SNVs where $0.3D < D < 3D$. Only sites in coding sequences of annotated genes (including genes annotated as hypothetical) were retained. SNVs that were absent in >0.25 of samples in a colony were removed. For colony-level estimates of SNV diversity, measures of coverage and allele frequencies for each retained SNV were combined across replicate samples. Filtered SNV counts in both sample-specific and colony-specific analyses were used to estimate the prevalence of SNVs over time, defined as the proportion of samples (specific to a given colony) where the minor allele comprises the majority of reads. As the reference allele is defined arbitrarily by the reference genome, the prevalence estimates to polarize each SNV based on the consensus across all time points of a given colony. Polarized alleles specific to each colony were then defined as the fraction of reads supporting the minor allele across time points. The coverage within some sample at a given time point, and the reads attributed to the major and minor frequency at $i$ allele, are referred to as $D_t$, $R_{it}$, and $A_{it}$, respectively. At any time point, $D_{it} \sim R_{it} + A_{it}$.

##### 0.6.3 Quantifying SNVs over time

To quantify SNVs within each MAG, as seen in Figure 4A and others, we adapted a metric of intermediate-frequency polymorphism similar to ones developed in [14, 18]. For each time point SNVs were retained if SNV coverage was $0.3\bar{D} < D < 3\bar{D}$. We then collected SNVs where $0.2D_{it} \leq A_{it} \leq 0.8D_{it}$ across 75% time points associated with the MAG × Colony pair. Sites that violated this requirement were kept but were not used in downstream analyses.

##### 0.6.4 Estimating core-genome diversity

To compare measures of variation between MAGs, we calculated several common statistical measures of genetic variation. Nucleotide diversity ($\pi$), which measures the average number of pairwise differences between any two sequences [21] was calculated from SNVs as follows:

$$\pi = \frac{\sum_{i<j} k_{ij}}{n(n-1)/2}$$

where $k_{ij}$ equals the number of nucleotide differences between the $i$th and $j$th sequences in the sample and the denominator represents the number of unique comparisons made between $n$ sequences

[21]. $\theta_w$, which is an alternative estimator of $\theta$, was calculated as follows where $S$ is equal to the total number of segregating sites (or SNVs):

$$\theta_w = \frac{S}{a}$$

where $a$ (a normalizing factor representing the sample size (n) is calculated from:

$$a = \sum_{i=1}^{n-1} \frac{1}{i}$$

Because both $\pi$ and $\theta_w$ are both estimators of the same parameter $\theta$, the expected difference between them should be 0 under the standard neutral model. We estimated the differences between $\pi$ and $\theta_w$ via *Tajima's D* [22] as follows:

$$D = \frac{\pi - \theta_w}{\sqrt{Var(\pi - \theta_w)}}$$

Finally, to evaluate short-term selective pressures within individual colonies, we calculated genic ratios of non-synonymous to synonymous polymorphism rates ($pN/pS$) [23] as below:

$$\frac{pN}{pS} = \frac{dN/nN_{ref}}{dS + 1)/nS_{ref}} \tag{14}$$

where $dN$ is calculated as the total number of non-synonymous alleles and $nN_{ref}$ is the total number of 0-fold degenerate sites in the reference gene. Similarly, $pS$ is calculated by dividing the total number of synonymous alleles ($dS$) by the total count of 4-fold degenerate sites in the reference gene ($nS_{ref}$). A smoothing pseudo-count of +1 was added to the $dS$ count in order to prevent undefined ratio when zero synonymous alleles were observed.

##### 0.6.5 Quantifying SNV frequency trajectories and differences between time points

To quantify changes in the genetic composition of individual MAGs, we searched for SNVs that underwent changes in allele frequency between any two time points. The type of change observed was defined as one of two classes. In the first class, allele frequency change between time points was simply defined was the difference in $A_{t1}$ and $A_{t2}$ for time points 1 and 2, respectively. In the second class, we adapted methods from Roodgar et al [18] to define SNVs that undergo full or partial "sweeps" due to large changes in allele frequency between time points. We use the term $\Delta RAF$ for variants satisfying these criteria and do not infer a selective sweep from the trajectory alone. However, as noted by Roodgar et al, estimating true allele frequency changes from spurious signals produced by finite sampling of reads in each time point is challenging. The uncertainty of allele frequency change increasingly decreases at higher coverages, but scanning for large allele frequency changes across many sites and measures of temporal distance can amplify error rates. Here, we report our adaptation of the approach developed in [18, 14] for detecting genetic differences between pairs of timepoints but suggest reading both for their intuition and insight.

For any pair of consecutively sampled timepoints, $t_1$ and $t_2$, we identified SNVs where $A_{it}$ transitioned from $< 20\%$ frequency in $t_1$ to $> 70\%$ in $t_2$. Under a null hypothesis of no allele frequency change, the probability of observing an event by chance through sampling noise is given here:

$$\begin{aligned} P_i(t_1,t_2) = & \underbrace{F(0.2\mathcal{D}_{it_1}|\mathcal{D}_{it_1},\bar{f})F(0.3\mathcal{D}_{it_2}|\mathcal{D}_{it_2},1-\bar{f})}_{\text{from } \leq 0.2 \text{ to } \geq 0.7} \\ & + \underbrace{F(0.2\mathcal{D}_{it_1}|\mathcal{D}_{it_1},1-\bar{f})F(0.3\mathcal{D}_{it_2}|\mathcal{D}_{it_2},\bar{f})}_{\text{from } \geq 0.8 \text{ to } \leq 0.3}, \end{aligned} \tag{15}$$

where $= (A_{it_1} + A_{it_2})/(D_{it_1} + D_{it_2})$ is the average allele frequency between any two timepoints. $F(k|n,p)$ is the cumulative distribution function for a binomial distribution with sample size $n$ and probability $p$ for success. For the null hypothesis, the expected number of differences is given by

$$n_{err}(t_1,t_2) = \sum_i P_i(t_1,t_2) \tag{16}$$

which can then be compared to the observed value $n_{obs}(t_1,t_2)$. In order to maximize statistical power to detect these large frequency changes, we excluded sites from both observed and expected counts if they met any of the following criteria:

1. **High false positive rate:** $P_i(t_1,t_2) > 10^{-3}$
2. **Insufficient polymorphism**: $A_i \leq 0.2$ in all timepoints.
3. **Variable copy number**: $\frac{D_{it}}{D_t}$ differed by more than a factor of 2 between timepoints $t_1$ and $t_2$

Based on the above definitions, the observe and expected SNV differences between all pairs of time points originating from a single colony was calculated. SNV frequency changes between time point pairs were considered if (i) the estimated false discovery rate $n_{err}(t_1,t_2)/n_{obs}(t_1,t_2) < 0.1$ and (ii) the Benjamini-Hochberg-corrected $P$-value for total number of observed changes is less than 0.05. When conditions were met, we marked all SNVs that were observed to change between this timepoint pair. As elaborated in [18], this approach leverages correlations between SNVs that change "together" to help overcome the high uncertainty associated with the frequency trajectory of a single SNV. This procedure was performed on all MAGs with at least 2 timepoints. For each MAG, we assigned temporally variable SNVs as the union of all SNVs identified in any significant timepoint pair. We note here, in line with [18], that this approach will miss many true allele frequency changes, and that our estimates of allele frequency changes should be treated as a lower bound in our experiment. In practice, allele frequency changes that do not meet our thresholds or increase slowly over multiple time points will not be captured. These methods would benefit from work that leverages shared frequency correlations across multiple timepoints, however developing these methods is beyond the scope of the current study.

Validation of allele frequency estimates with biological replciates Due to the nature of our study design, we sought to increase our confidence in allele frequency changes by leveraging observations across independent metagenomic replicates at each timepoint. For each significant SNV allele frequency change at the colony level, we identified timepoints with ≥2 replicate samples. Within each replicate and at the colony-level, we identified SNVs that were presented in all replicates for that timepoint and the SNVs present in the colony-polarized sample for that timepoint. We required shared SNVs to be present in ≥25% of timepoints. Colony-replicate comparisons that maintained ≥1000 shared, persistent SNVs were used to calculate the frequency within each replicate. Allele frequency values for each replicate were then compared to the pooled allele frequency value. For each persistent SNV in each timepoint, we used Pearson correlations between the replicate and colony-level allele frequency to create distributions of allele frequency differentials at the replicate level. In practice, this approach compared how well our pooled-colony allele frequency approximated the allele frequency observed in each biological replicate.

###### 0.6.6 Calling synonymous and nonsynonymous variants

In order to differentiate between SNVs contributing to synonymous or nonsynonymous diversity, we first identified all codon positions within genes containing SNVs. We then identified the relative position of each SNV within each codon, allowing us to compare the ancestral (e.g. the codon containing the polarized major allele) and derived (e.g. codon containing the polarized minor allele) codon sequence. Both codons were translated using a standard bacterial genetic table (Table 11). SNVs were assigned degeneracy based on the potential impact of the SNV on the codon-encoded amino acid, where 0D SNVs always resulted in an amino acid change and 4D SNVs never resulted in an amino acid change. Intermediate degeneracy scores (2D and 3D) were also classified. SNVs were then defined as synonymous or nonsynonymous if the SNV changed the amino acid encoded by the derived codon relative to the amino acid encoded by the ancestral codon. Mutations producing stop codons or codons that could not be unambiguously translated were excluded from downstream analyses.

##### 0.6.7 Estimating temporal Fst

To quantify the divergence within microbial populations over time, we calculated genic $F_{st}$ [24] between timepoints. For any given pairwise timepoint, $F_{st}$ was calculated as below:

$$F_{st} = \frac{H_T - H_S}{H_T} \tag{17}$$

$H_S$ describes the overall diversity at either timepoint ($T_1$ and $T_2$) and is defined as the mean heterozygosity for a given gene at polymorphic sites shared by both subpopulations. $H_S$ was then calculated as the unweighted average of the two timepoints: $H_S = (T_1 + T_2)/2$. $H_T$ describes the overall diversity of the combined subpopulations and is defined as the mean pooled heterozygosity across shared polymorphic sites in a given gene.

With the measures above, we calculated $F_{st}$ in one of two ways. 1) Month-to-month $F_{st}$ was calculated between consecutive timepoints (e.g. May to June, June to July). 2) Cumulative divergence was calculated between the initial timepoint (usually May) and all sequential months in order to measure cumulative genetic divergence.

##### 0.6.8 Estimating resident genomic backgrounds

To better distinguish evolutionary modifications of resident bacterial species from ecological mechanisms, such as strain replacement, we measured *marker SNVs* in each MAG in each colony. Our method is inspired by methods in [18], and has been used in previous studies to quantify sharing or transmission of bacterial strains across different hosts.

For each MAG $\times$ colony population, the first longitudinal observation criteria was designated $t_0$. High-frequency marker alleles ($> 0.9$) associated with this initial population were oriented such that the resident allele had frequency $f_{\text{resident}} = 1$ at baseline. Marker observations that were not callable at subsequent timepoints were treated as missing rather than as loss of the resident allele.

For each month $t$, resident-background turnover was defined as:

$$B_t = 1 - median(f_{resident}, t) \tag{18}$$

where $f_{resident,t}$ is the allele frequency of callable marker alleles. Values near zero indicate preservation of the genomic background observed at $t_0$, whereas larger values indicate progressively larger displacement of these marker alleles.

Populations were required to contain at least 20 marker al leles, at least 20 of these alleles at each point, and a minimum of 0.8 of these alleles available at each timepoint. We considered the initial background retained at any given month when:

$$B_t \leq 0.20 \tag{19}$$

For any timepoint transition from $t_1$ to $t_2$,

$$|\Delta B| = |B_{t2} - B_{t1}| \tag{20}$$

Rapid changes in allele frequency were considered ”resident-preserving” if:

$$B_{t2} \leq 0.20 \; and \; |\Delta B| \leq 0.10 \tag{21}$$

We note that these are operational thresholds and are not treated as definitions of discrete strains or haplotypes.

##### 0.6.9 Genome-wide $\Delta RAF$ and abundance calculations

We hypothesized that large changes in the resident genetic background should be accompanied by many simultaneous changes in allele frequency. Alternatively, thousands of simultaneous $\Delta RAF$ events could be inferred by large disruptions in strain composition, which result in the inflation of allele frequency shift counts. Estimating the breadth of these allele frequency changes alongside large disruptions in the genetic background would better distinguish $\Delta RAF$ events associated with large strain replacements versus those associated with highly localized adaptive events. We interpret broad shifts (high proportion of genome exhibiting $\Delta RAF$ events) as multi-locus or genome-wide shifts in allele frequency driven by large changes in the resident genetic background. In contrast, we interpret narrow shifts as those occurring across few genes or sites occurring *in situ* within a preserved resident genetic background. For each MAG x colony x consecutive timepoint, genome-wide $\Delta RAF$ was defined as:

$$S = \frac{N_{\Delta RAF}}{N_{trackable}} \tag{22}$$

where $N_{\Delta RAF}$ is the number of unique SNVs satisfying our $\Delta RAF$ criteria during a timepoint pair and $N_{trackable}$ is the number of unique SNVs within each of the timepoints. Duplicate records referring to the same SNV were collapsed before calculating either quantity. The magnitude of abundance change across a timepoint pair was therefore calculated as:

$$A = |log_2 \frac{p_{t2}}{p_{t1}} \tag{23}$$

where $p_t$ is the corrected relative abundance a focal population. Timepoint pairs with $A \leq 1$ were described as accumulating less than a twofold change in abundance.

Associations between shift breadth, $|\Delta B|$, and abundance change. $A$, were quantified using Spearman correlations. Confidence interval were obtained by cluster bootstrap, which resamples complete MAG x colony longitudinal trajectories with replacement so that repeated observations from the same microbial population were not treated as statistically independent. The same boostrap procedure was used to estimate the confidence intervals for the proportion of background-preserving $\Delta RAF$ timepoint pairs.

##### 0.6.10 Null model of purifying selection for$\pi_N/\pi_S$ and model selection

To better understand how purifying selection structures genetic variation observed within and across bee-associated bacterial populations, we examined the relationship between nonsynonymous nucleotide diversity relative to synonymous nucleotide diversity, $\pi_N/\pi_S$, and synonymous nucleotide diversity $\pi_S$. This analysis was motivated by recent models developed to study analogous questions within the human microbiome [14] where the efficiency of purifying selections is dependent on the timescale over which deleterious mutations are purged from the population. In the original model, Garud et al observed a rapid decline in $d_N/d_S$ with increasing $d_S$, where $d_S$ can be interpreted as a proxy for the effective age of mutations or the depth of divergence among distinct, host-specific conspecific strains [14]. Here, we adapted this logic to include nucleotide diversity and used $\pi_S$ as a proxy for standing natural diversity and the genealogical. While conceptually similar, we note that our approach has limitations relative to the original $d_N/d_S$ approach in that we are unable to directly estimate the fraction of deleterious mutations. Our decision to use this approach was motivated by the fact that few MAGs were associated with $> 1$ colony, which prevented the use of $d_N/d_S$ estimates.

For each MAG $\times$ colony $\times$ month sample, we estimated $\pi$ at 0-fold degenerate sites and 4-fold degenerate sites. We define $\pi_N$ as nucleotide diversity at 0-fold generate codon positions, which we treat as nonsynonymous diversity, and $\pi_S$ as nucleotide diversity at 4-fold degenerate codon positions, which we treat as synonymous diversity. For some gene $i$, we denote $\pi_{N,i}$ and $\pi_{S,i}$ as gene-level nonsynonymous and synonymous diversity, respectively. We further denote $L_{0D,i}$ and $L_{4D,i}$ as the effective number of callable 0D and 4D sites in that gene. Both $L_{0D,i}$ and $L_{4D,i}$ allow us to control for the difference in opportunity for 0D and 4D diversity between genes. For each MAG $m$, colony $c$, and month $t$, we aggregated gene-level diversity using weighted means based on the effective length of each gene, where:

$$\pi_N^{(m,c,t)} = \frac{\sum_i \pi_{N,i}^{(m,c,t)} L_{0D,i}}{\sum_i L_{0D,i}} \tag{24}$$

$$\pi_S^{(m,c,t)} = \frac{\sum_i \pi_{S,i}^{(m,c,t)} L_{4D,i}}{\sum_i L_{4D,i}} \tag{25}$$

The nonsynonymous diversity ratio for a given sample was defined as:

$$R^{(m,c,t)} = \frac{\pi_N^{(m,c,t)}}{\pi_S^{(m,c,t)}} \tag{26}$$

Samples were retained for model fitting only if $\pi_N^{(m,c,t)}$ and $\pi_S^{(m,c,t)}$ were finite and greater than zero. We additionally required each MAG × colony *month* sample to contain a minimum number of genes contributing diversity estimates to reduce the influence of sparse observations. Unless stated otherwise, all analyses were performed on aggregated MAG × colony × month observations. We note that gene-level analyses were also aggregated for pathway-specific analyses.

Our use of $\pi_S$ follows from coalescent theory, where neutral nucleotide diversity reflects the the mutation rate and mean coalescent time of a lineage. Under the assumption that 4D sites are apprixmately neutral with respect to fitness, low $\pi_S$ corresponds to relatively shallow genealogies or low standing neutral diversity whereas higher values of $\pi_S$ correspond to deeper genealogies, larger effective population sizes, or both [25, 21]. We therefore treated $\pi_S$ as an axis along with which the efficacy of purifying selection is expected to vary. At low $\pi_S$, nonsynonymous variants may be younger on average and therefore expected to be less efficiently removed from the population. This results in an elevated $\pi_N/\pi_S$. In contrast, when $\pi_S$ is high, purifying seleciton has had a longer effective timescale with which to remove deleterious nonsynonymous variants. This is expected to lower $\pi_N/\pi_S$. We emphasize that $\pi_S$ is not a direct estimate of mutation age, effective population size, or coalescent time. Variation in mutation rate, recombination, linkage between sites, population structure, strain composition, migration, and recent demographic change can all contribute to variation in measures of $\pi_S$. We therefore interpret $\pi_S$ as a descriptive proxy for both neutral diversity and genealogical depth.

Further, we note that a concern when analyzing $\pi_N/\pi_S$ as a function of $\pi_S$ is that $\pi_S$ appears as both axes (as a measurement on the x-axis and as the denominator in the y-axis). Shared sampling noise in $\pi_S$ can generate a negative relationship between $R$ and $\pi_S$, even in the absence of any biological effect. In order to decouple measurements of $\pi_S$, we implemented a gene-level splitting procedure that is analogous to the Poisson thinning approaches used in divergence-based analyses [14]. For each MAG $\times$ colony $\times$ month sample, genes were randomly assigned to one of two groups. We then computed two independent estimates of $\pi_S$:

$$\pi_{S,1}^{(m,c,t)} = \frac{\sum_{i \in G_1} \pi_{S,i}^{(m,c,t)} L_{4D,i}}{\sum_{i \in G_1} L_{4D,i}} \tag{27}$$

and

$$\pi_{S,2}^{(m,c,t)} = \frac{\sum_{i \in G_2} \pi_{S,i}^{(m,c,t)} L_{4D,i}}{\sum_{i \in G_2} L_{4D,i}} \tag{28}$$

We then defined a decoupled response based on:

$$R_{\text{decoupled}}^{(m,c,t)} = \frac{\pi_N^{(m,c,t)}}{\pi_{S,1}^{(m,c,t)}} \tag{29}$$

Model fitting was performed using $R_{\text{decoupled}}$ as the denominator in $\pi_N/\pi_S$ and $\pi_{S,2}$ for the predictor variable (x-axis). Because $\pi_{S,2}$ is estimated from a different set of genes than the denominator $\pi_{S,1}$, the persistence of any relationship does not originate due to sampling noise shared between the two variables.

We then compared several functional descriptions of the relationship between $\pi_N/\pi_S$ and $\pi_S$. First, we fit a mechanistic model adapted from the time-dependent purifying-selection seen in Garud et al [14, 17].

$$R_{\text{decoupled}} = (1 - f_d) + f_d \frac{1 - \exp(-k\pi_{S,2})}{k\pi_{S,2}} \tag{30}$$

This models a decline in $\pi_N/\pi_S$ with increasing $\pi_S$ where $f_d$ is the fraction of deleterious mutations and $k$ is a selection coefficient. In the limit of small $\pi_{S,2}$, both $(1 - \exp(-k\pi_{S,2}) / (k\pi_{S,2})$ approaches and the expected ratio of $\pi_N/\pi_S$ approach 1. In the limit of large $\pi_{S,2}$, this term approaches 0, and the expected $\pi_N/\pi_S$ ratio approaches the lower asymptote $(1 - f_d)$. In this model, $f_d$ controls the amplitude of the decline in $\pi_N/\pi_S$ while $k$ controls the scale of $\pi_S$ over which the decline occurs.

Because our analysis is based on polymorphism-based estimates rather than divergence, we do not interpret $f_d$ or $k$ as direct estimates of the fraction of deleterious mutations or selection coefficients, as seen in the original implementation of the model in Garud et al. Instead, these values as factors influencing curve shape and interpret this model as a phenomenological null model that describes the amplitude and curvature of the empirical relationship between nonsynonymous diversity relative to synonymous diversity and standing neutral diversity. This differs from the original divergence-based model in two important ways. First, the original model was applied to divergence calculated among host-specific conspecific strains. Second, in the divergence-based analyses, $d_S$ can be interpret more directly as a proxy for divergence time or mutation age, whereas in our analysis $\pi_S$ reflects standing neutral diversity and is affected by demographic and ecological processes. Despite these differences, this model provides a useful baseline for asking whether $\pi_N/\pi_S$ declines with increasing neutral diversity in a manner that is consistent with an increasing efficiency of purifying selection in removing deleterious variants.

We fit both an unweighted and weighted version of this model, where the unweighted model treats each MAG $\times$ colony $\times$ month observation equally. The weighted model accounts for unequal variance in the data arising from unequal numbers of callable synonymous sites among genes. Each observation was weighted by the total effective number of callable synonymous sites used to estimate the two split synonymous diversity values:

$$w^{(m,c,t)} = L_{4D,1}^{(m,c,t)} + L_{4D,2}^{(m,c,t)}. \tag{31}$$

The weighted nonlinear least-squares model minimized the following weighted residual sum of squares:

$$\text{WRSS} = \sum_{m,c,t} \left( e_w^{(m,c,t)} \right)^2. \tag{32}$$

This is analogous to the process of minimizing residuals after multiplying both the observed and fitted values by $\sqrt{w^{(m,c,t)}}$. This weighting gives greater influence to MAG $\times$ colony $\times$ month observations when more synonymous sites are available, and therefore more accurate estimates of synonymous diversity. In addition to this model, we fit several empirical curves, including a Hill-type model:

$$R_{\text{decoupled}} = c \; + \; \frac{1-c}{1 + \left(\frac{\pi_{S,2}}{x_0}\right)^h}, \tag{33}$$

where $c$ is the lower asymptote, $x_0$ is the value of $\pi_{S,2}$ at which the curve transitions between high and low $\pi_N/\pi_S$, and $h$ controls how sharply this transition occurs. This model was included as a flexible empirical description of the same qualitative pattern without requiring the specific functional form motivated by the model drawn from [14]. We also fit a generalized additive model to describe the relationship nonparametrically:

$$\log_{10} R = s\left(\log_{10}(\pi_S)\right) + \epsilon \tag{34}$$

where $s(\cdot)$ is a penalized smooth function estimated from the data and $\epsilon$ is the residual error term. The GAM was included to test whether the data supported a monotonic decline in $\pi_N/\pi_S$ with increasing $\pi_S$ without requiring a specific parametric curve. Because both $\pi_N/\pi_S$ and $\pi_S$ span several orders of magnitude, the GAM was fit on log-transformed values, matching the scale used for visualization and reducing the influence of extreme values.

Model evaluations were made through comparisons of AIC. Agreement among the different model fits was interpreted as evidence that the observed decline in $\pi_N/\pi_S$ with increasing $\pi_S$ was not dependent on a single parameter. Together, these models were used as a descriptive null model for the observed genome-wide relationship between nonsynonymous diversity relative to synonymous diversity and standing neutral diversity.

##### 0.6.11 Spatial covariance of synonymous allele frequency change with focal $\Delta RAF$ events

To test whether rapid nonsynonymous changes in allele frequency were associated with change in local synonymous alleles, we assigned $0D\ \Delta RAF$ as focal sites and quantified allele frequency change at nearby synonymous sites (4-fold degenerate or 4D SNVs). Distances between SNVs were calculated only for SNV pairs on the same assembled contig to control for differences in assembly and gene organization across MAGs. To reduce heterogentity among genes, the primary analysis included only 4D SNVs located on genes separate from the gene containing the focal $0D\ \Delta RAF$.

For each focal event and nearby 4D SNV $i$, we calculated the absolute change in allele frequency over the same MAG x colony x timepoint,

$$|\Delta AF_{4D,i}| = |f_{i,t2} - f_{i,t1}| \tag{35}$$

Because genome-wide allele frequency dynamics exhibited high variation among populations and sampling intervals, 4D allele frequency change was normalized to a transition-wide background, where transitions are defined as any consecutive timepoint pair (e.g. May-June, June-July, etc).

$$E_i = |\Delta AF_{4D,i}| - median(|\Delta AF_{4D}|)_{MAG\ X\ Colony\ X\ Interval)} \tag{36}$$

Nearby 4D SNVs were grouped into predefined distance bins. To prevent focal events containing many nearby SNVs from strongly influencing the analysis, excess synonymous change was first summarized within each focal event and distance bin before being summarized across all focal events.

As a control for spatial covariance unrelated to rapid focal shifts, we repeated the above analysis around matched 0D loci that did not satisfy our $\Delta RAF$ critiera. Control loci were matched within the relevant population and timepoint pair to account for bias and analyzing using the same distance bins and normalization procedure. This analysis was used to test for biases specific to $0D\Delta RAF$ events.

##### 0.6.12 Temporal covariance of synonymous allele frequency change with focal $\Delta RAF$ events

Spatial covariance alone cannot distinguish whether a specific genomic region is enriched for genetic variation relative to a region that becomes unusually dynamic during a focal $0D$ $\Delta RAF$. To test for this, we aggregated the behavior of focal sites over across all consecutive sampling intervals.

For each focal $0D$ $\Delta RAF$, the local region was defined as all callable 4D SNVs located within 5 KB of the focal site, on the same assembled contigs, and in genes different from where the 0D focal site was observed. For each available transition $t$, local synonymous allele frequency change was summarized simply as:

$$L_{r,t} = median_{i \epsilon r} |\Delta AF_{4D,i,t}| \tag{37}$$

where $r$ denotes the focal genomic region. Excess synonymous change for region $r$ was then defined as:

$$T_r = L_{r,t_{focal}} - median_{t \neq t_{focal}} L_{r,t} \tag{38}$$

Positive values indicate that the local region experienced greater synonymous allele-frequency movement during a focal $0D$ $\Delta RAF$ interval than during other observed transitions. To visualize these differences, transitions were indexed relative to the focal interval ($t = 0$), and local synonymous allele frequency change was centered relative to each focal region's expected change in synonymous allele frequency before aggregating across focal events.

Signifcance was assessed using a within-region permutation test. For each focal genomic region, one of its observed transitions was randomly designated as a pseudo-focal interval, while the region, population, set of local 4D SNVs, and observed temporal variation were held constant. The excess diversity statistic ($T_r$) was recalculated across all regions for each permutation. This procedure essentially asks whether the observed $0D$ $\Delta RAF$ interval is more locally dynamic than some randomly selected interval from the same genomic region.

We then compared the observed median excess in synonymous allele frequency change, the proportion of regions with positive $T_r$, and the proportion for which the focal interval was the most dynamic observed interval against corresponding permutation distributions. Probabilities were calculated as $(r+1)/(B+1)$, where $B = 1000$.

##### 0.6.13 Protein clustering across MAG core genomes

To identify proteins in each MAG's core genome that exhibited sequence similarity similarity, we clustered protein sequences from retained genes using MMseqs2 [19]. Proteins were clustered using *mmseqs easy-cluster –min-seq-id 0.5 -c 0.85 –cov-mode 0*, which produced 59,399 sequence-defiend protein families. Of these, 18,468 were represented in at least two unique MAGs. Functional annotations derived from PFAM and GO were assigned to protein families for interpreting gene function and were not used to define homology or similarity.

##### 0.6.14 Identification of recurrent rapid allele frequency changes at nonsynonymous sites

To identity allelic frequency changes of potentially adaptive significance, we prioritized $\Delta RAF$ occurring in 0D sites, as these are likely to have the largest impact on organism fitness via modification of the amino acid. Using these $\Delta RAF$ events, we searched for evidence for parallelism. In previous studies [26, 27, 28], parallelism is typically defined as recurrent mutations in different subpopulations of the same species – typically occurring in different hosts. Our approach differs from these existing studies due to two primary difficulties: 1) The lack of shared MAGs across individual colonies dramatically reduced the sequence space with which to search for parallel 0D $\Delta RAF$. 2) In order to control for HGT in SNV calling, we removed any genes that exhibited significant similarity when found between two different MAGs. Due to these challenges, we instead mapped 0D $\Delta RAF$ to clustered protein families generated in the previous step. $\Delta RAF$ were first aggregated at the gene level followed by association at the level of the protein family cluster. We then searched for examples of recurrent 0D $\Delta RAF$ within this clustered protein family. A recurrent family represents independent nonsynonymous $\Delta RAF$ within genes that share the same protein cluster but are associated with $\geq 2$ unique MAGs. We considered 0D $\Delta RAF$ as putatively concurrent when they satisfied the following criteria:

1) The 0D $\Delta RAF$ occurs in $\geq 2$ unique MAGs

2) The 0D $\Delta RAF$ occurs in MAGs belonging to two different genera

3) The 0D $\Delta RAF$ occurs in MAGs belonging to distinct colonies

4) The 0D $\Delta RAF$ occurs in two genes found within the same clustered protein family

Recurrent 0D $\Delta RAF$ that met these criteria were retained. As an additional layer of scrutiny, we further filtered recurrent 0D $\Delta RAF$ by temporal concurrence. We defined temporal concurrence as recurrent 0D $\Delta RAF$ that met all existing criteria while also occurring during the same month pair (e.g. between May and June). Exact month-to-month intervals, rather than broad seasonal categories, were used to define temporal concurrence.

##### 0.6.15 Permutation testing for recurrent $\Delta RAF$

In order to test whether temporally concurrent recurrence exceeded expectations based on random $\Delta RAF$ occurrence, we constructed a null distribution in which the observed number of 0D-swept genes within each MAG x colony x time-interval ($N$) was kept constant. For each permutation, we sampled $N$ genes without replacement from genes within the corresponding MAG x colony population. Sampling probability was kept proportional to the effective number of 0D sites within each gene. This procedure allows us to preserve population-specific $\Delta RAF$ opportunity while accounting for both the variable number of $\Delta RAF$ occurring over time as well as the variation in the number of 0D sites across genes. Randomized genes were then mapped to their associated clustered protein family, and we then counted the number of distinct protein families that satisfied the same recurrence criteria seen above. The permutation statistic was the number of protein-family x sampling-interval combinations that satisfied this definition. This test was performed 5,000 times, and we calculated a one-sided probability as $\frac{r+1}{N+1}$, where r is the number of permutations producing a statistic equal to or greater than the observed value.

##### 0.6.16 Protein family residue localization and structural analysis of recurrent amino acid variants

Protein sequences from retained core genes were clustered with MMseqs2 (–min-seq-id 0.5 -c 0.85 –cov-mode 0) 19, and members of recurrent candidate families were aligned with MAFFT. Positions of $0D\delta RAF$ were mapped from the original gene sequence to the corresponding MMseqs2 protein and subsequently through the family alignment onto a representative medoid sequence. Original amino acid coordinates were obtained from codon_order_in_gene. Mappings were retained only when the original protein-to-MMseqs2-member and family-alignment coordinate transformations could be validated through alignment. Residue identities displayed on representative structures therefore correspond to the medoid protein and do not necessarily represent the ancestral or derived amino-acid states observed in individual populations.

Within-protein localization was evaluated by permuting residue labels across MAGs. Mapped positions were randomized among valid medoid residues while preserving the number of events contributed by each MAG. Multiple events within a MAG were collapsed to their median swept position, and the span among independent MAG-level positions was compared with the permutation distribution.

The exact 249-aa medoid sequence was structurally predicted with ColabFold v1.6.2 using AlphaFold2-ptm. The highest-ranked model had pTM = 0.90 and mean pLDDT = 94.65; pLDDT values at positions 33, 39, 52 and 77 were 98.12, 98.69, 98.31 and 89.44, respectively. Pairwise C and minimum heavy-atom distances were calculated from predicted coordinates. Positions 33, 39 and 52 formed a compact structural region with a maximum pairwise C distance of 13.24 Å. This geometry was stable across all five ranked predictions (cluster diameter, 13.20–13.29 Å).

The conserved Walker A motif (GSSGSGKS, residues 43–50) and ABC signature motif (LSGGQ, residues 145–149) were identified from the medoid sequence. Minimum heavy-atom distances were calculated between mapped positions the and Walker A motif .Position 52 was 2.87 Å from the motif in the highest-ranked model and 2.82–2.89 Å across all five predictions.

In order to more deeply explore the structural context of these mutations, we superimposed the medoid model onto the experimentally determined ATP-bound ABC transporter nucleotide-binding-domain dimer 1L2T using UCSF ChimeraX MatchMaker. The structures aligned with an RMSD of 1.10 Å across 163 pruned atom pairs (2.46 Å across all 219 aligned pairs). ATP coordinates from 1L2T were retained solely as an experimental structural reference and were not modeled as ligand-bound coordinates for the focal protein. Minimum heavy-atom distances from medoid positions 33, 39, 52 and 77 to template ATP were 16.04, 12.61, 4.79 and 20.65 Å, respectively; position 52 was nearest the phosphate region of ATP (4.79 Å).

##### 0.6.17 0D seasonal $\Delta RAF$ rate

To test whether nonsynonymous $\Delta RAF$ rates differed across seasons, we counted the number of 0D $\Delta RAF$ events occurring in a gene in a season for each gene-season combination. We then fit GLMMs of the form:

$$sweeps_{gene,season} \sim Poisson(\lambda) \tag{39}$$

where

$$log(\lambda) = \beta_0 + \beta_{season} + log(length\ of\ nonsynonymous\ sites) \quad (40)$$

The log of nonsynonymous site counts was included as an offset term to control for the unequal opportunity of 0D $\Delta RAF$ across genes. This allows us to model the rate of nonsynonymous $\Delta RAF$ events per site rather than raw $\Delta RAF$ counts. We included genus identity as an additional covariate to account for taxon-specific differences in $\Delta RAF$ dynamics across the colonies.

Model coefficients were used to obtain differences in $\Delta RAF$ rates among seasons relative to the baseline season (Fall). Gene-level $\Delta RAF$ rates were then summarized as the number of nonsynonymous $\Delta RAF$ events per million nonsynonymous sites. Model-based seasonal $\Delta RAF$ rate estimates and confidence intervals were derived from the Poisson regression models.

###### 0.6.18 Testing for the effect of drift among allele shifts in time series

To test whether allele frequency changes observed in Figure 5 and 6 are consistent with a simplified model of genetic drift, we implemented a simplified variant of the statistical model developed by Roodgar et al [18]. Rather than fit a full hidden Markov model to each individual SNV trajectories, we instead estimated the effective population size ($N_\mathrm{e}$) of each species $\times$ host combination using allele-frequency variation at four-fold degenerate (4D) SNVs. These sites are used to estimate the expected frequency change at a given site.

The core intuition of this approach is that the size of allele-frequency fluctuations generated by genetic drift scales inversely with $N_\mathrm{e}$. By estimating $N_\mathrm{e}$ from the observed frequency variation over the course of the experiment, we can derive an estimation of the degree of allele-frequency change that is expected from drift and sequencing noise alone. We then test the observed $\Delta RAF$ trajectories against this expectation.

For each MAG $\times$ colony pair, we identified putatively neutral SNVs using three increasingly permissive site classes: synonymous sites (4D), all non-0D sites (nonsynonymous), and all polymorphic sites. Sites that were previously identified with an allelic $\Delta RAF$ were excluded from this set. The observed allele-frequency change that we observed at each class of SNVs as computed at sequential timepoint pairs:

$$\Delta f = f_{t+1} - f_t \tag{41}$$

where $f_t$ denotes the observed allele frequency at the initial timepoint. We further calculated the number of generations that occurred within each timepoint ($\tau$), which is based on the number of days separating the timepoints and an assumed bacterial growth rate of 12 generations per day. The number of generations per day is assumed based a doubling time of 2 hours, which we have observed from bee-associated microbial cultures growing across various media. The real doubling time within honey bees is likely slower.

Following the Wright–Fisher diffusion approximation, the expected variance in allele-frequency change generated by neutral drift over a time interval $\tau$ generations scales with both allele frequency and effective population size. Rather than explicitly fitting the diffusion process, we used the variance expected under this model to estimate $N_{\mathrm{e}}$ and construct a neutral null expectation for temporal allele-frequency change.

We summarized allele-frequency variation across all neutral intervals using three quantities:

$$S_1 = \sum_i f_i(1 - f_i)\tau_i, \tag{42}$$

$$S_2 = \sum_i f_i(1 - f_i)\left(\frac{1}{D_i} + \frac{1}{D_{i+1}}\right), \tag{43}$$

$$S_3 = \sum_i (\Delta f_i)^2, \tag{44}$$

where $D_i$ and $D_{i+1}$ denote sequencing coverage at the beginning and end of each timepoint pair. Intuitively, $S_1$ represents the cumulative amount of heterozygosity exposed to drift through time, $S_2$ represents the variance expected from sequencing and sampling noise, and $S_3$ represents the total observed variance in allele-frequency change.

where $D_i$ and $D_{i+1}$ denote sequencing coverage at the beginning and end of each timepoint pair. Intuitively, $S_1$ represents the amount of expected heteroyzogisty that is exposed to drift through time, $S_2$ represents the expected amount of sequencing noise, and $S_3$ represents the total observed allele-frequency variance.

Effective population size was then estimated using a moment-based estimator

$$\hat{N}_e = \frac{S_1}{2(S_3 - S_2)} \tag{45}$$

which we use to define the excess variance in allele-frequency change beyond what is expected from sequencing noise. This variance is attributed to genetic drift. We retained estimates only when the observed variance exceeded sampling variance ($S_3 > S_2$) and a minimum number of timepoint pairs were available for a given MAG $\times$ colony pair. This procedure was performed independently for 4D sites, non-0D sites, and all sites. In order to maximize the conservativeness of the null model, we retained the minimum estimate among these three estimations ($\mathrm{N}_{\mathrm{e,min}}$) for each MAG. Because the expected excess variance due to genetic drift increases with decreasing effective population size, the minimum estimate maximizes the amount of allele-frequency change that can be attributed to neutral processes and makes subsequent tests more conservative.

For each $\Delta RAF$, we calculated the expected variance contributed by sequencing noise,

$$V_{\text{sample}} = f(1-f)\left(\frac{1}{D_t} + \frac{1}{D_{t+1}}\right), \tag{46}$$

and the expected variance contributed by genetic drift,

$$V_{\text{drift}} = f(1-f)\left(1 - e^{-\tau/(2N_e)}\right), \tag{47}$$

which corresponds to the variance accumulated under the Wright–Fisher diffusion approximation over $\tau$ generations. For each timepoint pair, we calculated the sampling variance,

$$V_{\text{sample}} = f(1-f)\left(\frac{1}{D_t} + \frac{1}{D_{t+1}}\right). \tag{48}$$

and the drift variance

$$V_{\text{drift}} = f(1-f)\left(1 - e^{-\tau/(2N_e)}\right). \tag{49}$$

The total variance for each timepoint pair was then simply:

$$V_{\text{tot}} = V_{\text{sample}} + V_{\text{drift}}. \tag{50}$$

Observed allele-frequency changes throughout sampling were standardized as:

$$Z = \frac{\Delta f}{\sqrt{V_{\text{tot}}}}. \tag{51}$$

such that values near zero are consistent with neutral expectations. Large deviations from zero indicate greater allele-frequency shifts than expected from drift and sequencing noise alone.

Each longitudinal allele trajectory was then summarized by:

$$Z_{max} = \max_t |Z_t| \tag{52}$$

Large values of $Z_{max}$ are indicative that $\geq 1$ observed timepoint transition deviated strongly from an expectation based on drift-plus-sampling. The simulated null distribution was built by drawing values directly from a zero-mean normal distribution, scaled by each interval's specific standard deviation. Under genetic drift and sampling noise, the standardized change in allele frequency across any time interval ($Z = \frac{\Delta f}{\sqrt{V_{tot}}}$) is expected to have a mean of 0 and variance of 1. The $|Z| = 1.96$ represents a pointwise 95% normal boundary and is not treated as a significant threshold. Transitions that fall outside of this distribution (e.g. $> 2$ are interpreted as being significantly different from the null expectation.
